# Combined transcriptomic, connectivity, and activity profiling of the medial amygdala using highly amplified multiplexed *in situ* hybridization (hamFISH)

**DOI:** 10.1101/2024.09.14.613026

**Authors:** Mathew D. Edwards, Ziwei Yin, Risa Sueda, Alina Gubanova, Chang S. Xu, Virág Lakner, Megan Murchie, Chi-Yu Lee, Kristal Ng, Karolina Farrell, Rupert Faraway, Subham Ganguly, Elina Jacobs, Bogdan Bintu, Yoh Isogai

## Abstract

*In situ* transcriptomic technologies provide a promising avenue to link gene expression, connectivity, and physiological properties of neural cell types. Commercialized methods that allow the detection of hundreds of genes *in situ*, however, are expensive and therefore typically used for generating unimodal reference data rather than for resource-intensive multimodal analyses. A major bottleneck is the lack of a routine means to efficiently generate cell type data. Here, we have developed hamFISH (highly amplified multiplexed fluorescence *in situ* hybridization), which enables the sequential detection of 32 genes using multiplexed branched DNA amplification. We used hamFISH to profile the projection, activity, and transcriptomic diversity of the medial amygdala (MeA), a critical node for innate social and defensive behaviors in mice. In total, we profiled 643,834 cells and classified neurons into 16 inhibitory and 10 excitatory types, many of which were found to be spatially clustered. We then examined the organization of outputs of these cells and activation profiles during different social contexts. Therefore, by facilitating multiplexed detection of single molecule RNAs, hamFISH provides a streamlined and versatile platform for multimodal profiling of specific brain nuclei.

## Introduction

A key milestone for the study of neural circuits is to delineate the composition and connectivity of each brain area that controls specific behaviors. Systematic identification of the individual components of the circuits *i.e.*, neuronal cell types, has seen a dramatic breakthrough in recent years^1^. This includes not only the transcriptomic and proteomic compositions of individual cells but also their morphology, biophysical properties, and connectivity with other brain areas. Understanding these basic parameters is essential to map information flow and to gain insights into potential computations within circuits^2^. In addition, each circuit component should be characterized in the context of the patterns of activity during different behaviors.

The multimodal characterization of cell types, however, remains technically challenging. While *in situ* transcriptomic methods allow cell type assignment in the native tissue context and are amenable to combining with other neural readouts, existing methods including commercial platforms, suffer from two major limitations for wide adoption. First, *in situ* transcriptomics experiments are resource intensive. While they provide vital spatial information for transcriptomic cell types, they have not been used as routinely as one might need for multimodal experiments. Second, most platforms use decoding of barcoded signals, requiring a complex analysis pipeline. One way to address these issues is to limit *in situ* transcriptomics experiments to specific brain areas and use a small number of marker genes to perform the profiling. There are several existing platforms^3, 4^ one can use to perform these experiments efficiently for up to 24 genes with sequential gene readout. Higher number of genes, however, remains more challenging due to complex requirements on probe designs and extended experimental timelines. To address these points, we have developed hamFISH (<u>h</u>ighly <u>a</u>mplified <u>m</u>ultiplexed fluorescence *in situ* <u>h</u>ybridization), a method that uses branched DNA amplification on a well-established smFISH platform with gene-specific bridge readout probes to sequentially detect gene expression. Additionally, the use of an optimized tissue preparation protocol to detect mRNA transcripts in brain tissue sensitively and flexibly makes hamFISH more amenable for routine use. We then validated the robustness of hamFISH in the medial amygdala (MeA), a critical hub for processing social information that receives prominent pheromonal input^5, 6^. Transcriptomic cell types in the MeA were recently defined by single-cell sequencing^1, 7, 8^. We therefore decided to use these datasets as a starting point to map these cells in a multimodal fashion, *i.e.*, their spatial localization, projection targets, and activation profiles during distinct innate social behaviors.

Using a panel of 31 genes, we profiled 643,834 cells and classified MeA neurons into 16 inhibitory and 10 excitatory neuronal cell types, many of which have distinct spatial patterns. Furthermore, hamFISH has allowed us to identify a rare inhibitory cell type (*Ndnf^+^*) localized in the molecular layer of the MeA. We then combined hamFISH with MeA output mapping and found cell-type biases in projections to the medial preoptic area (MPOA), bed nucleus of the stria terminalis (BNST), and the ventrolateral portion of the ventral medial hypothalamus (VMHvl). Finally, by combining hamFISH with *c-fos* activity mapping upon interactions with conspecific males, females, or pups, we found that instead of specific MeA transcriptomic types being activated by unique social stimuli, most of these types were combinatorially activated, generating unique MeA population activities. Our *in situ* profiling of the MeA, therefore, provides a proof-of-principle example of overlaying transcriptomic types, projection, and activity in a behaviorally relevant manner and demonstrates the usefulness of hamFISH in multiplexed *in situ* gene expression profiling.

## Results

### Robust detection of RNAs in the brain using highly amplified multiplexed FISH

To achieve robust transcript detection under the high autofluorescence observed in mouse brain tissues, we first tested branched DNA amplification^9^ with single-molecule FISH. We designed preamplifier and amplifier oligonucleotides that contain repetitive sequences that together multimerize to form branched DNA. In the original scheme (highly amplified version 0), the preamplifiers were designed to bind common non-complimentary overhangs (20 nt) on RNA-specific encoding probes (**Figure 1A**). Preamplifiers also include 9x binding sites (19 nt) for amplifier oligonucleotides which themselves contain 10x binding sites (18 nt) for fluorophore-conjugated readout probes (all sequences are compiled in **Table S1**). We found that this method increased signal intensity by 2.7-fold (807 ± 20 A.U. vs 2157 ± 63 A.U.). However, we observed a high number of punctate signals (39.2 ± 4.4 puncta/10 µm^2^) in the extracellular space (**Figure 1B,C** and **Figure 1 – figure supplement 1A**). To test if these extra-somatic signals are true signals or represent non-specific probe binding events, we designed probe sets that recognize two different regions of *Parvalbumin* (*Pvalb*) mRNA (n=12 probes for each), each with a distinct fluorescence readout (**Figure 1 – figure supplement 1B**). Extra-somatic signals had an overlap rate of 6.2 ± 1.0%, significantly lower than perinuclear signals in *Pvalb^+^* cells (39.1 ± 2.4 %), suggesting that the bulk of these spots may be non-specific signals (**Figure 1 – figure supplement 1C, D**). To alleviate this issue, we modified the probe design so that preamplifier binding sequences were split into two 14-nt half sites on adjacent encoding probes (highly amplified version 1.0) similar to splitFISH^10^. Indeed, we found that the extra-somatic signal decreased when using this design (17.4 ± 1.4 puncta/10 µm^2^, **Figure 1B** and **Figure 1 – figure supplement 1A**). Furthermore, there was an increase in the overlap rate of perinuclear *Pvalb^+^* spots (46.5 ± 2.5%) relative to our first design suggesting an improvement in the fidelity of signal detection (**Figure 1 – figure supplement 1D**). Compared with non-amplified smFISH, this branched DNA amplification scheme increased smFISH signal by 3.8-fold (807 ± 20 A.U. vs 3072 ± 50 A.U., **Figure 1C**).

**Figure 1.**
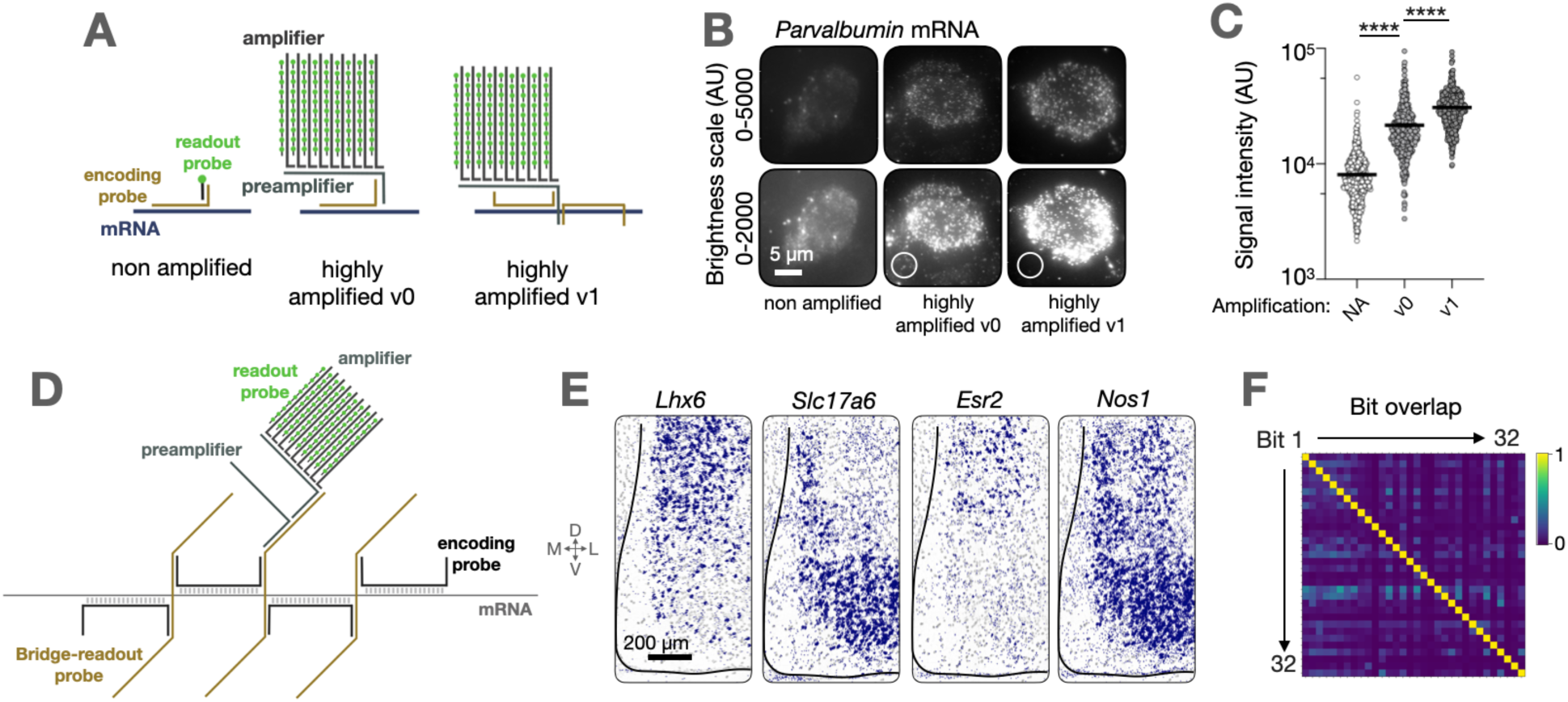
Highly amplified, multiplexable smFISH in the medial amygdala. **A.** Schematic of smFISH readout systems. **B.** Representative images from Parvalbumin smFISH comparing amplification methods. Circles indicate examples of extracellular regions with non-specific signals. **C.** Quantification of smFISH signal intensity from different amplification methods, n = 500 signals (50 from 10 cells) for each condition. Signal intensity is background subtracted (mean of 50 background intensities subtracted from transcript signal). p > 0.0001, unpaired T-test. **D.** Schematic of final hamFISH readout system incorporating bridge-readout probes and highly amplified v1 design. **E.** Representative images known MeA marker gene transcripts in the MeA, detected using the hamFISH scheme shown in D. **F.** Proportion of bits that give overlapping single molecule localized signals in a sequential smFISH experiment where all 32 bits were used to develop a different gene.

Separately, we developed a system with preamplifiers and amplifiers containing 32x repeats (see materials and methods for details) to enable the reliable detection of low-abundance transcripts. These amplifiers, termed version 2.0, increased the single molecule signals by 12.3-fold compared with the non-amplified signal (807 ± 20 A.U. vs 9849 ± 172 A.U.), and 3.2-fold over our highly amplified version 1.0 smFISH method (**Figure 1 – figure supplement 2A**). Moreover, we performed rolling circle amplification (RCA) mediated detection of *Pvalb* transcripts^11^. Although the probe chemistries of these methods are significantly different, we found that the signal intensity from the version 2.0 method and RCA-based signals are comparable (**Figure 1 – Figure Supplement 2B**). These data therefore confirm that branched DNA amplification enables tuneable amplification of single molecule FISH signals. To enable multiplexing with the branched DNA trees, we designed additional amplifiers and tested them using different encoding probes. Curiously, not all designed amplifiers produced specific signals with high dynamic range (n=13/32). We therefore replaced the amplifier sets by screening new sequences and selecting only those that produced a high signal-to-noise ratio in a more standardized fashion using *Sst* mRNA as a common target (see materials and methods for details).

To utilize encoding probes amplified from oligopools, which offer a significant cost advantage over individually synthesized probes, we modified our original scheme to include bridge-readout probes (**Figure 1D**). These gene-specific probes create an intermediate layer that links encoding and preamplifier probes. Importantly, while oligopool probes are synthesized in a pool and fixed in composition, bridge-readout probes are individually synthesized and can be customized for each experiment. This scheme therefore allowed us to flexibly select a subset of genes for readout without resynthesizing oligopool libraries.

The amplifiers were then tested for their orthogonality. We used an encoding probe library in which each gene is coupled to a unique amplifier set, confirming that the incorporation of the bridge-readout probe was successful at detecting marker genes in the MeA (**Figure 1E,Figure 1 – figure supplement 3**). Upon imaging signals from all the amplifiers sequentially, we performed single-molecule localization and examined signal overlaps. We then excluded those that showed cross-talk, resulting in an orthogonal set of 32 amplifiers (**Figure 1F**). Our amplified smFISH design therefore enables transcript detection in brain tissue that is compatible with multiplexed spatial profiling methods.

### Application of hamFISH for profiling a subnucleus of the amygdala

Compared to *in situ* transcriptomics methods using barcoding at a single molecule level, non-barcoded, sequential readout of multiplexed FISH signals does not necessitate the precise alignments of single molecule signals from imaging rounds. In addition, amplified signals using hamFISH enabled us to speed up the imaging time by using a lower magnification objective (40x as opposed to 60x). Multiplexing by fluorescent readouts using AlexaFluor-488, Cyanine 3 (Cy3) and Cyanine 5 (Cy5) further reduced imaging time.

This scalability makes sequential hamFISH an attractive method to be coupled with other physiological and connectivity readouts, where increased numbers of samples are inevitable due to the lower throughput nature of physiological and neural tracing methods. As an important case study, we applied hamFISH for cell type profiling linking gene expression, connectivity, and activity in the medial amygdala (MeA). The MeA is a critical hub for the neural circuits underlying innate social and defensive behaviors in mammals. While the behavioral roles of the MeA have been well established and specific MeA subpopulations have been linked to the execution of specific innate behaviors^12, 13^, circuit-level mechanisms of how each MeA cell drives specific physiological and behavioral effects remain incompletely characterized. Addressing cell type diversity in multimodal fashion is an important step in delineating the circuit basis of how these neurons control a repertoire of behavior that includes mating, aggression, and parenting.

To choose a panel of genes that captures cell type diversity in the MeA, we first re-clustered previously published scRNA-seq MeA data^8^ and found 26 clusters in cells that are *Slc32a1* (inhibitory) or *Slc17a6* (encoding Vglut2 and therefore excitatory) positive (**Figure 2 – figure supplement 1A, Table S2**). Eight of these clusters we suspected of either being located outside the MeA, based on Allen Brain Atlas *in situ* data (clusters 6, 11, 14), or expressed ubiquitously throughout the MeA and surrounding areas (clusters 4, 5, 20, 21, 26). In addition, we found three clusters that did not have specific marker genes that were differentially expressed or seemed to be a mix of excitatory and inhibitory cells (clusters 16,17,25), leaving 15 MeA cell clusters. Therefore, we assembled a gene panel consisting of marker genes for these scRNA-seq clusters and those used in previous studies. We finalized 31 genes for subsequent hamFISH experiments (**Table S3**).

We then used sequential hamFISH to profile cells along the full anterior-posterior axis of the MeA (at 100 µm interval over 1 mm). Initially, experiments were performed on 13 animals with a total of 163,925 cells profiled (mean of 12,607 ± 351 per animal). Of these, 40.3 ± 0.5% were inhibitory and 39.0 ± 1.4% excitatory, as defined by the expression of *Slc32a1* and *Slc17a6* respectively. The remaining cells were made up of either cell types that are localized outside of the MeA and did not express either *Slc32a1* or *Slc17a6* (8.6 ± 0.7%) or were MeA cells whose identity could not be established due to the absence of any markers other than *Slc32a1* or *Slc17a6*. Nevertheless, cells in which the expression of three or more marker genes is detected made up ∼80% of the cells in the MeA. This is consistent with the scRNA-seq data showing that approximately 20% of the MeA cells are glia. Our 31-gene panel can therefore be used to define most of the neurons in the MeA.

We then clustered our hamFISH data for *de novo* discovery of cell types and identified 16 inhibitory (i1-i16) and 10 excitatory (e1-e10) neuronal classes, along with 4 classes of cells that expressed neither *Slc32a1* nor *Slc17a6* (n1-n4) (**Figure 2A-C, Figure 2 – figure supplement 2A-B**). The clusters were consistent between animals and sexes (**Figure 2 – figure supplement 2C**). To determine cluster reproducibility between different profiling methods, we correlated the mean expression of our gene panel markers in our hamFISH clusters with clusters from Chen *et al.* (2019) scRNA-seq data (**Figure 2 – figure supplement 1B**). We found reasonable correspondence between the datasets. For example, scRNA-seq clusters 1, 7 and 8 contained *Slc17a6*, *Adcyap1*, *Nos1*, *Penk,* and *Tac1* positive cells, which correspond to hamFISH clusters e1-e6. The excitatory classes fall approximately into 3 classes: e1-6, e7-9 and e10 when compared with sequencing data. There were also hamFISH clusters that were not found in the scRNA-seq dataset. We attribute them to the clusters with very low number of cells (i3, i4) or low expression of marker genes that may not be reliably detected by scRNA-sequencing (i7-i9). In more recently published scRNA-seq data^1^, however, we find these clusters except i3, perhaps due to the deeper sequencing coverage.

**Figure 2.**
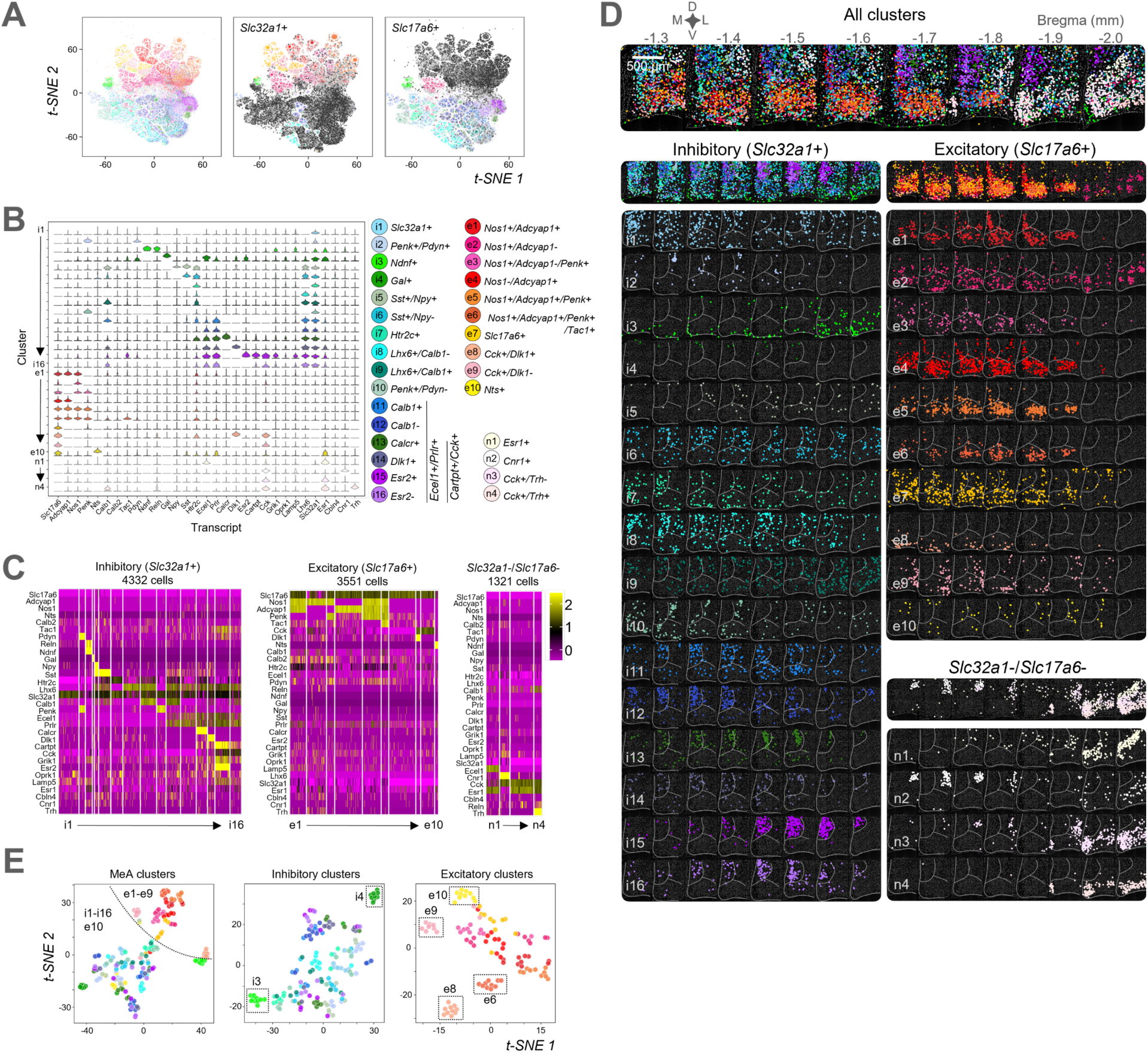
The MeA is comprised of distinct molecular cell types that occupy specific spatial locations. **A.** Two-dimensional t-distributed stochastic neighbor embedding (t-SNE) plots based on differentially expression of 31 genes for n = 163,925 cells from 12 animals, colored by cluster. Those cells positive for either *Slc32a1* (inhibitory) or *Slc17a6* (excitatory) are highlighted in the right two panels. **B.** Marker gene expression distributions within each cluster, represented by violin plots for cells shown in A. **C.** Heatmap representation of the 30 identified clusters from one animal (male). Note gene order differences between plots to aid with visualization of clusters. **D.** The spatial position of every cell from each cluster from one animal (male) across the MeA. White lines represent MeA subdivision structure drawn manually based on DAPI morphology and cluster positions. **E.** t-SNE plots based on spatial features of each cluster. MeA clusters (left) have been separated into inhibitory (middle) and excitatory (right) classes. Color scheme the same as indicated in B, separable clusters are indicated with boxes.

We then proceeded to analyze the spatial distributions of these clusters. Consistent with previous work^14^, most inhibitory classes are localized in dorsal and ventral subdivisions of the MeA, whereas excitatory neurons occupy primarily the ventral MeA (**Figure 2D, Figure 2 – figure supplement 2D**, **Figure 1E**). We found, for example, that the posterodorsal MeA (MeApd) contained six unique *Ecel1^+^Prlr^+^* inhibitory clusters (i11-i16) with specific spatial patterns. For example, among the clusters marked by the expression of *Cartpt* and *Cck,* i15 (*Esr2^+^*) and i16 (*Esr2^-^*) occupy the most posterior part of the MeApd, and these clusters are flanked laterally by i13 (*Calcr^+^*) neurons and intermixed with the sparser i14 (*Dlk1^+^*) population (**Figure 2B**). Rostral to these posterior MeApd populations are two subtypes that do not express either *Cartpt* or *Cck* but are positive or negative for *Calb1* (i11 and i12, respectively). Excitatory classes also show specific spatial structures. For example, *Cck*^+^ excitatory cells can be split into a *Dlk1*^+^ population (e8) that solely occupies the most ventral part of the anterior MeA (MeAa), and a *Dlk1*^-^ population (e9), which sits more laterally. Other striking spatial classes of MeA neural subtypes include sparse inhibitory *Sst*-expressing populations (i5, i6) that span the entire MeA in a salt-and-pepper pattern and *Ndnf*^+^ inhibitory cells (i3) that line the medial border with the optic tract, or ventral surface of the MeA.

Our MeA hamFISH data also include cell populations outside the MeA based on the discontinuity of marker expression. For example, *Penk^+^Pdyn^+^* (i2) cells primarily occupy a location lateral to the MeA, possibly the central amygdala (CeA), with a smaller proportion that appeared to localize within the MeA. There are four clusters located in the CeA (n2) and posteromedial cortical amygdala (PMCo, n1, n3 and n4) which are *Slc32a1*^-^ and *Slc17a6*^-^ (n1-n4). These are putatively *Slc17a7*^+^ excitatory cells based on scRNA-seq and Allen Brain Atlas *in situ* data (mouse.brain-map.org). hamFISH therefore enabled the spatial mapping of at least 16 inhibitory and 10 excitatory neuron types.

Moreover, to compare differences in the spatial distribution of each cluster and quantify reproducibility between experiments, we coarsely aligned sections from different experiments and clustered cell classes using metrics that describe the spatial components of each cluster (**Figure 2E**). These were: mean x (medial-lateral axis) position, standard deviation (SD) of x, mean y (dorsal-ventral axis), SD of y, mean z (anterior-posterior axis), SD of z, total number of cells, distance to the molecular layer (the input layer of the MeA), and the distribution of clusters that harbor nearest neighboring cell to each neuron in consideration. The clustering revealed that inhibitory and excitatory classes generally have different spatial properties (**Figure 2E**, left), although the salt-and-pepper, sparse nature of e10 (*Nts*^+^) cells is more similar to inhibitory cells than other excitatory classes. Within the inhibitory classes, i3 (*Ndnf*^+^) and i4 (*Gal*^+^) are separated into distinct clusters, consistent with a more restricted spatial pattern compared with other inhibitory clusters (**Figure 1E**, middle). Within the excitatory classes, e6, e8, e9 and e10 were separable from other clusters, consistent with more ventral positions (e6 and e8), lateral position (e9) and sparse salt-and-pepper position (e10) compared with other excitatory cells (**Figure 1E**, right). These data therefore show that many of the transcriptomic cell clusters have distinct spatial patterns. By combining our amplified smFISH design with a panel of 31 genes imaged sequentially, our data provide a comprehensive *in situ* map of the major transcriptomic neural clusters in the MeA.

### hamFISH uncovers a rare population of inhibitory neurons in the MeA

Among the three classes of MeA GABAergic neurons that have been characterized *in vitro*, only one type, which is localized to the molecular layer, harbors short axons with fast-spiking characteristics, and has therefore been proposed to be a local interneuron^15, 16^. Strikingly, our *in situ* map of MeA neuron types identified a single class of cells preferentially localized to the MeA molecular layer (i3), and we decided to characterize this population further. These cells express *Ndnf* and account for 95% of inhibitory neurons localized there. Notably, these cells are rare, comprising only 1.8% of neurons in the MeA, and we failed to identify this subtype in the reference scRNA-seq data. These cells also express *Slc32a1* and *Reln*, but not *Npy* or *Nos1*, and therefore resemble a specific class of cortical L1 interneuron that has been linked to gain modulation^17^, plasticity, and cross-modal integration^18^. As the acetylcholine receptor a7 (encoded by *Chrna7)* is also expressed in GABAergic L1 neurons^19^ we tested if the remaining population of GABAergic cells in the molecular layer correspond to a different type. Unexpectedly, we found that 74.5% of *Ndnf^+^* cells in the molecular layer of the MeA express also *Chrna7* (**Figure 2 – supplemental figure 3A**), indicating that there is further heterogeneity within the *Ndnf^+^* population.

To further characterize the connectivity of MeA*^Ndnf^* neurons, we mapped their inputs and outputs. Using *Ndnf-Cre* animals in which Cre-dependent AAV harboring synaptophysin-EGFP and tdTomato was injected into the MeA, we identified the axonal projections of these neurons (**Figure 2 – figure supplement 4A)**. We found GFP signal in four major brain areas: MeA, AOB, cortical amygdala (olfactory cortex) and BNST, indicating that, unlike L1 *Ndnf^+^* neurons, MeA*^Ndnf^* neurons harbor both local and long-range axonal arbors (**Figure 2 – figure supplement 4B,C)**. MeA i3 neurons especially send a strong long-range input to the AOB, including mitral and granular cell layers, and to the BNST. Conversely, mapping of the inputs to i3 neurons by rabies tracing showed that they receive inputs from various brain areas including olfactory areas, cortex, and hippocampal formation (**Figure 2 – Figure supplement 4D-F**). Notably, a similar retrograde tracing experiment using *Sst*-expressing neurons in the MeA as starter cells showed these neurons have different input structures. In particular, MeA*^Ndnf^* neurons receive more inputs from olfactory areas and less inputs from the dorsal striatum compared to MeA*^Sst^* neurons. Altogether, these anatomical data establish that MeA^Ndnf^ neurons represent a unique neural subtype embedded in the circuit processing both main olfactory and vomeronasal inputs.

### The projection specificity of transcriptomic subtypes of MeA neurons

Combining gene expression profiling of neural tissue with other biological readouts such as connectivity is vital to understand how neural circuits function. Our simple method to profile transcriptomic types *in situ* facilitates further multimodal interrogation of these cells. MeA neurons send projections to both proximal and distal targets^14, 15, 20^. We sought to determine the projection targets of specific cell classes by combining hamFISH with multicolor retrograde tracing. We injected fluorescently labeled cholera toxin subunit b (CTB) into three limbic regions known to receive strong MeA inputs and play important roles in social behaviors: the medial preoptic area (MPOA), the posterior medial bed nucleus of stria terminalis (BNST), and the ventrolateral subdivision of ventromedial hypothalamus (VMHvl) **(Figure 3A, Figure 3 – figure supplement 1A**). Initial quantification of labeled cells in the MeA revealed differences in the distribution of projecting cells along the anterior-posterior axis (**Figure 3B, Figure 3 – figure supplement 2A**). Specifically, VMHvl-projecting neurons were enriched in the anterior over posterior, with a significant decrease at bregma -1.7 mm. In contrast, both BNST- and MPOA-projecting cells were distributed evenly through the anterior-posterior axis. We also compared the total number of CTB-positive cells per target area in the MeA and found that BNST-projecting cells were nearly 2-fold more abundant than MPOA- and VMHvl-projecting cells (**Figure 3 – figure supplement 2B**). Furthermore, triple color retrograde tracing in single animals allowed us to identify a minority of double- or triple-labeled cells, indicating that some MeA neurons target multiple brain regions (**Figure 3 - figure supplement 2C, D**). Approximately 75% of neurons were singly labeled, which is unlikely due to inefficient labeling as the rate of triple BNST-injected positive neurons was near 100% (**Figure 3 – figure supplement 1B**). These observations suggest specific connectivity of each MeA cell to downstream targets.

**Figure 3.**
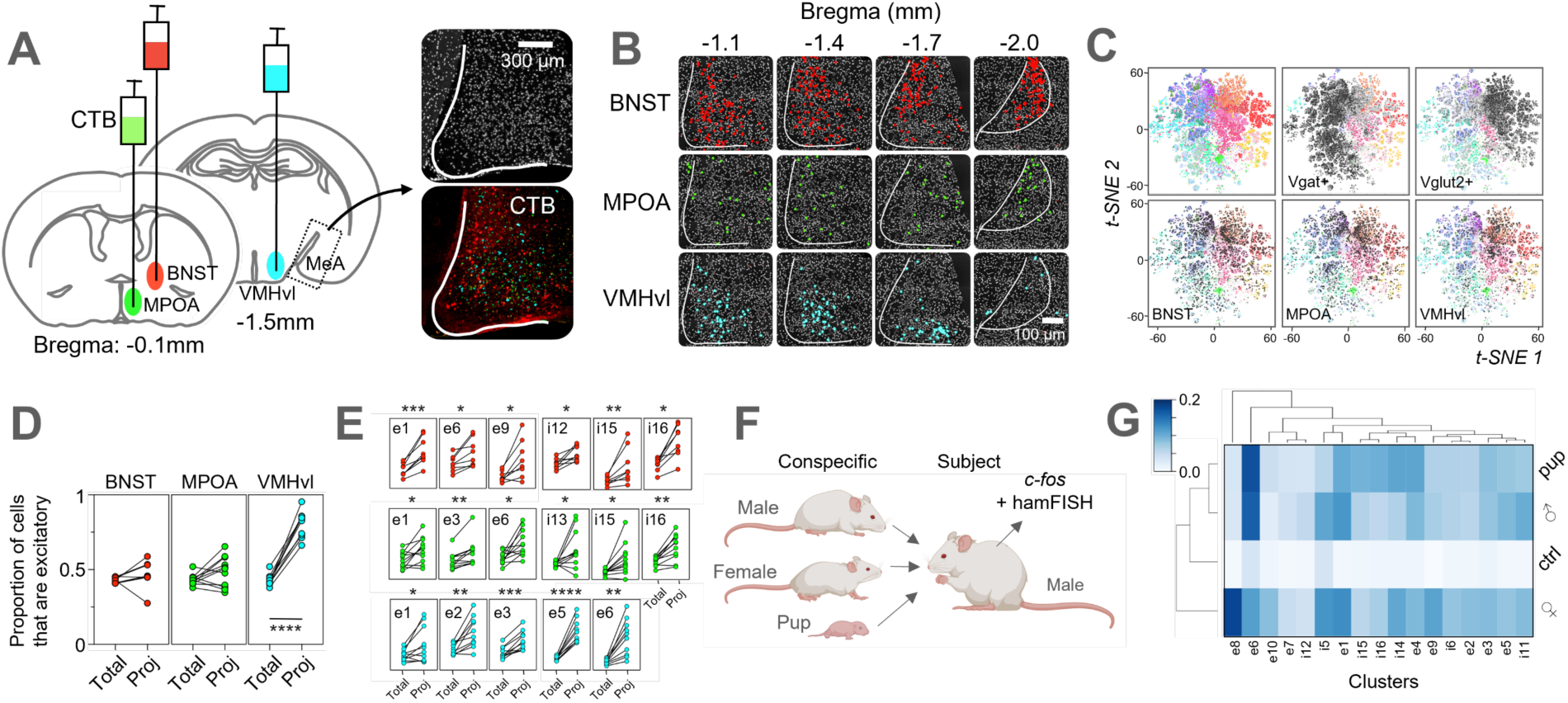
Specific MeA cell types are associated with projection targets and are activated differentially to social stimuli. **A.** Schematic of CTB-injection scheme along with a representative MeA image from triple-injected animals. Note that injection sites of the 3 different CTB-conjugated fluorophores were swapped around in different animals, but for the purpose of this figure all subsequent data adhere to the color scheme green = MPOA-projecting cells, red = BNST-projecting cells, blue = VMHvl-projecting cells. **B.** Representative position of CTB-labelled projection neurons in a single animal injected in all 3 target regions. **C.** *t*-SNE plots. Shaded cells in top row represent either inhibitory or excitatory cells (left panel is *t*-SNE without shading). Bottom row shading indicates cells labelled with CTB injected into either the BNST, MPOA or VMHvl. n = 263,474 cells from 13 animals, coloured by cluster. **D.** Comparison of the proportion of all cells that are excitatory vs projection neurons labelled by CTB that are excitatory. For VMHvl *p* < 0.0001, paired T-test. **E.** Comparison of the proportion of labeled CTB positive cells for each cluster. Paired T-tests. **F.** Schematic of activity mapping workflow. **G.** Hierarchical clustering of mean *c-fos* expression per cluster in exposed mice. Scale bar represents proportion of cells in cluster positive for *c-fos*.

We then classified the labeled cells into the transcriptomic classes as defined above. Using hamFISH we assigned 263,474 CTB-labeled cells from 13 animals to neuronal cell types (**Figure 3C, Figure 3 – figure supplement 3**). These experiments uncovered two major insights into MeA projection neuron specificity. First, at the mesoscale level, we found a significant enrichment in VMHvl-projecting neurons that were excitatory (**Figure 3D**). In fact, of all VMHvl-projecting cells, 80 ± 9 % were excitatory in contrast to 43 ± 4 % of all neurons assigned. BNST and MPOA-projecting cells were instead made up of a similar proportion of excitatory cells as non-projecting cells (46.7 ± 9 % for BNST-projecting cells, and 49 ± 10 % for MPOA-projecting cells). Second, we found CTB labeling in most of the transcriptomic clusters, indicating most projection neurons project to all three areas examined. There was, however, enrichment of labeling in specific clusters for certain projection targets (**Figure 3E, Figure 3 – figure supplement 2E**). We found that VMHvl-single-projecting neurons, for example, were over-represented in the excitatory clusters defined by the marker genes *Adcyap1*, *Nos1* and *Penk* (e.g., e2-e6), when compared with BNST and MPOA-single projecting cells. Broadly, MPOA and BNST-projecting neurons showed similar cell type enrichment to each other despite our findings that BNST/MPOA-double-projecting neurons were relatively low in abundance (6-13%) (**Figure 3 – figure supplement 2D**). Although MPOA and BNST-projecting neurons were assigned to excitatory cell types in the ventral MeAa, a greater enrichment could be seen in inhibitory *Ecel1*^+^*Prlr*^+^*Lhx6^+^* MeApd clusters (clusters i8, i11-i16) when compared with VMHvl-projecting cells. Interestingly, BNST-projecting cells also showed enrichment in three excitatory clusters (e1, e4 and e8) when compared with MPOA-projecting cells, with e8 (*Cck^+^Dlk^+^*) cells also enriched compared with VMHvl-projecting cells (**Figure 3 – figure supplement 2E**). In addition, we identified several subtypes that are enriched in cells sending projections to multiple targets (**Figure 3 – figure supplement 2F,G**). For example, the cluster i16 was more abundant in double-projecting BNST/VMHvl and MPOA/VMHvl cells compared with single VMHvl-projecting cells. Conversely, clusters e3 and e6 were enriched in double-projecting BNST/VMHvl and BNST/MPOA cells compared with single BNST-projecting cells. These results therefore uncover a mixed but biased projection profile of individual transcriptomic MeA populations.

### Mixed selectivity of the activation patterns of MeA transcriptomic types upon innate social behaviors

We next tested the response of MeA neurons to social stimuli to determine the input organization to each MeA cluster. We prepared MeA sections from the animals exposed to different conspecifics in a freely behaving fashion. For this experiment, we also included probes against the immediate early gene *c-fos,* which has been commonly used to survey activity in this brain area^14, 21^. Virgin male mice in their home cage were presented with either another male, a female, or pups, so that neuronal activation could be correlated with aggression, mating, and infanticide, respectively (**Figure 3F**). We found that while many of the transcriptomic classes can be commonly activated during social behaviors, the pattern of activation appears to be stereotyped and different among different behavioral contexts (**Figure 3G** and **Figure 3 – figure supplement 3A**). We found only a few clusters selectively activated during specific social behavior (**Figure 3 – figure supplement 3B**). The excitatory cluster e6 (*Tac1^+^*) was more highly activated in both aggressive (14.4 ± 1.2 % of e6 cells were *c-fos^+^*) and infanticidal (15.0 ± 2.2 %) behaviors compared with reproductive behavior (10.1 ± 0.7 %, p<0.01 vs male, p<0.001 vs pup, paired t-Test), whereas e8 (*Dlk1^+^*) cells showed the opposite preference, being more highly activated during mating behavior (25.4 ± 3.0 % of e8 cells were *c-fos^+^* when exposed to a female conspecific compared with 2.8 ± 1.1 % in male exposed samples (p<0.0001), and 5.0 ± 2.3 % in pup exposed samples (p<0.05)). Activation of inhibitory clusters was similar during different behaviors, although i14 and i16 were more activated during aggressive behavior than infanticide. Taken together, these results suggest that different social stimuli activate the ensemble of transcriptomically distinct MeA neurons in unique patterns.

## Discussion

### Development of highly amplified multiplexed FISH as a routine method for multiplexed transcript detection

Various *in situ* methods have been developed to capture the transcriptomic complexity of neurons while preserving spatial information, e.g., MERFISH^22–24^, split-FISH^10^, seqFISH^25, 26^, STARmap^11^, coppaFISH^27^, Hyb-ISS^28^, EASI-FISH^3^ and BARseq2^29^, each offering unprecedented opportunities for understanding cell type heterogeneity within tissues. Furthermore, commercial platforms, *e.g.* MERSCOPE (Vizgen), Xenium (10x Genomics) and CosMx (NanoString) facilitate *in situ* transcriptomic studies using many (>100) genes, however, the high costs of these platforms preclude them from being used for routine multi-modal survey of cell types. In contrast, platforms such as coppaFISH^27^, EASI-FISH^3^, SABER^4^, and RNAscope^30^ allow cell type analysis with fewer genes (fewer than 100 but more than 5) and have been more amenable for multimodal analysis. These methods, however, pose challenges in implementing from scratch when commercial systems are not available.

Branched DNA amplification has been available for decades and is proven to be a robust platform for signal amplification in tissues^9^. Indeed, our initial tests using two complementary probe sets for a single gene confirm the detection of gene transcripts with rare dropouts. However, it remains challenging to extend this method beyond a dozen amplifier sets. We tested nearly 100 sequences to select specific amplifiers with high signal-to-noise and validated their orthogonality. This resulted in the development of 32 orthogonal amplifier modules. Our split encoding probe design is a feature that can be used in other methods such as splitFISH^10^ and HCRv3^31^ to ensure a low incidence of false positives. In addition, we designed and synthesized two different lengths of amplifiers to provide differential signal strength. Long amplifiers were used to read out *c-fos* to ensure the high dynamic range of expression of this gene. Short amplifiers (v1) produced sufficient signals for many of our target transcripts. In contrast, long amplifiers (v2) produced more signals, which some bled-through into unintended fluorescence channels under our filter setting.

Furthermore, the use of oligo pools to prepare probes rather than individually synthesized oligonucleotides that are commonly used for sequential detection of transcripts, helped reduce experimental costs. Different probe libraries can be independently amplified from single oligo pools that contain as many as 10,000 oligonucleotides. With the ‘bridge-readout’ scheme, one can further narrow down genes that should be visualized to optimize the marker gene sets. We also demonstrated that our probe scheme could be combined with other amplification systems such as hybridization chain reaction (HCR)^31, 32^ by incorporating HCR initiator sequences into bridge-readout probes. In addition, the imaging of each round was faster than HCR-based methods^3, 33^ which require stripping and polymerization steps. We estimate that the start-up reagent cost of hamFISH using 32x multiplexing is 7,000 GBP as of writing, which can accommodate approximately 140 experiments assuming that each sample covers 10 mm^2^ tissue. The system is versatile so that one could choose to multiplex less, which further reduces the cost of the experiments. Taken together, these developments should democratize *in situ* cell type identification in individual laboratories.

### Towards multimodal in situ map of MeA transcriptomic cell types

Previous behavior studies elucidated the functional specialization of unique MeA populations in specific social behaviors, e.g., MeApd GABAergic neurons are involved in aggression while glutamatergic neurons in MeApv are involved in self-grooming^12^. In contrast, despite single-cell RNA sequencing of MeA neurons uncovering further transcriptomic heterogeneity in the MeA^7, 8, 34^, the circuit role as well as behavior and physiological roles of most of these clusters remain uncharacterized. We therefore surmised that a constructive direction would be to overlay the anatomical and activity information on these transcriptomics datasets.

By using hamFISH as a workhorse method for *in situ* transcriptomics to survey over 600,000 cells, we learned that this focused approach complements unbiased sc/snRNA-sequencing screens. Our data revealed further subtypes that were missing from this dataset when compared to amygdala cell types found in scRNA-seq data available at the time of our data collection^8^, likely due to low abundance (i3) or those (i8, i9, i15) that are defined by genes with low expression levels (e.g., *Lhx6*, *Esr2*). Notably, we designed and collected the *in situ* data before new sets of data using 10x v3 platforms became available^1, 7^. Still, we have since found evidence of all these cell types in scRNA-sequencing data except for i3.

The mapping of MeA transcriptomic types uncovered a striking spatial organization of transcriptomic types within MeA subregions. The neuronal types that occupy specific locations include e8 and e9, which are excitatory ventral *Cck*^+^ cells, as well as inhibitory i13, i15, and i16 cells which express *Cartpt*, *Prlr*, *Cck* and *Ecel* that mark the posterodorsal MeA. We also found that some clusters exhibit a salt-and-pepper distribution within a specific region, which might add further heterogeneity in a given population. Moreover, we also identified a rare cell type localized in the input layer of the MeA. Notably, *Ndnf^+^* inhibitory neurons (i3) were not detected in the previous scRNA-seq data. Furthermore, we found expression of *Chrna7* in a proportion of *Ndnf^+^* cells, implying at least 2 different subtypes of cell make up the i3 population.

Recent studies identified two major developmental lineages of MeA neurons, *Dbx1*-derived and *Foxp2*-neurons^13, 35, 36^. Although *Dbx1* is not expressed in the adults and *Foxp2* was not included in our gene panel, the Allen ABC atlas^1^ shows that *Foxp2* is broadly expressed across several of our hamFISH clusters, for example *Sst^+^/Npy^-^* (i6), *Htr2c* (i7), *Calb1* (i9), and seems to overlap with the *Esr2^-^* MeApd population (i16). Therefore, given the roles of *Foxp2* neurons in aggression^13^, it would be interesting to study the role of each of the *Foxp2* subpopulations during conspecific male-male contact. Moreover, our projection mapping experiments uncovered that excitatory MeAa cells project to the VMHvl (**Figure 3 – figure supplement 2G**), an area that has been identified to play an important role in the experience-dependent escalation of aggressive behavior^37^.

Whether mammalian pheromones activate dedicated neural circuits as observed in insect systems remains unresolved. Indeed, ESP1 and ESP22 have been shown to activate *c-fos* within MeA neurons in stereotyped patterns, some of which appear to mediate the action these pheromones in the control of lordosis and mating^38, 39^. In contrast, *in vivo* physiological recordings consistently found population coding within this area^40, 41^. To this end, we found no molecularly defined clusters uniquely activated during specific behavioral outputs, with a notable exception of e8. Importantly, the activation pattern of each cluster as a MeA ensemble appears unique in different social contexts, providing a potential molecular link to the neural representation of social information in the MeA^40, 41^. Our experiments also found the robust activation of glutamatergic MeA neurons during social behaviors, perhaps consistent with the activation of ventral MeA by single pheromonal compounds^38, 39^, but their circuit and behavioral roles need to be uncovered by specifically manipulating these populations rather than using pan-glutamatergic cell markers, the activation of which produces self-grooming^12^. A prominent future direction therefore is to study a specific population of neurons based on molecular identity and projection targets, which would enable a systematic investigation of connections in this circuit.

### Pitfalls of the study

A major pitfall of hamFISH is that this method is currently limited to thin section samples (<20 µm) due to the large molecular size of amplifiers, which creates a barrier to diffusion. HCR provides better scalability to thicker tissues in this respect. In addition, hamFISH currently allows only 32 genes to be sequentially read out, thus this method may prove inadequate for profiling at a deeper transcriptomic level. For example, our study failed to identify sexually dimorphic cell types that had been previously reported^7^ indicating that our gene selection provides limited resolution. Additionally, profiling of multiple brain areas simultaneously would likely require a custom gene panel for each area. Future investigations therefore could use multiplexing strategies^42^ to profile the functional properties of MeA cells at their finest (*i.e.*, “cluster”) level in the MeA.

## Materials and Methods

### Animals and behavioral assays

All animal experiments were performed under the UK Animals (Scientific Procedures) Act 1986 (P108E8CF8) following local ethical procedures. Adult wild-type CD1 and C57BL/6J (Charles River, UK) mice aged 8-12 weeks were used in this study and kept on a 12h:12h light/dark cycle with *ad libitum* access to food and water. Only CD1 mice had tissue collected to be used for hamFISH experiments. Prior to behavioral assays for *c-fos* measurements, animals were individually housed for 7 days and then habituated in a testing room under dim red lighting for 3 hours before behavioral assays were performed (at 1-2h after the onset of the dark cycle). Animals were sacrificed 30 min after exposure to a conspecific before tissue was harvested. Behavioral assays that involved exposure to conspecifics were video recorded to ensure the selection of tissue from mice that displayed the appropriate behavioral responses (see below) for subsequent hamFISH experiments. The specific details of the behavioral experiments are described below.

*Infanticidal behavioral assays* – Pups were generated by mating CD1 males with C57BL/6 females. Infanticidal males were exposed to two freshly euthanized pups (P2-4) and tissue was selected for subsequent hamFISH assays if evidence of pup-directed aggression was confirmed (e.g., biting and eating).

*Inter-male aggression assays* – One male mouse (intruder) was taken from his cage and placed in the cage of another male mouse (resident). Tissue was used from the resident mouse if evidence of fighting was confirmed. This included bouts of biting and tail-rattling.

*Mating assays* – Male mice were introduced to the cage of a female mouse. Tissue was used from these mice if evidence of sexual behavior such as anogenital sniffing, mounting, and intromission was observed. The female oestrus cycle timepoint was confirmed by vaginal swab.

### Probe and bridge-readout design

Encoding probes were constructed by amplifying oligonucleotide sequences from complex oligo pools (synthesized by Twist Bioscience), as described below. To design the gene-specific sequences, we first used the BLAST program as 30 nt probe candidates against the mouse RefSeq dataset, then discarded sequences with hits with length 14 nt or more. Using R scripts, pairs of 30 nt were then selected in a manner that best optimized the number of consecutive probes along a transcript with a target of 90 sequences per gene, although some genes did not reach this due to sequence space. The reverse complement of the chosen sequences was then flanked with bridge-readout sites designed in the following way.

To design the flanking bridge-readout sites, a set of orthogonal 25mer DNA barcodes^42^ were expanded to 28mer sequences with the addition of random nucleotides, before a further BLAST search to exclude off-target hits. Only those where each half of the sequence had a similar T_m_ were chosen. Selected bridge-readout sequences were then split into 14mer half sites and placed on either side of the gene-specific sequences described above. Sequences were tested as described below. For the bridge-readout probes themselves, the complementary sequences of the 28mer bridge-readout sequences were conjugated to four preamplifier binding sites (two sites 5′ and two sites 3′) and synthesized as Ultramers by IDT. Flanking the encoding probe combined sequences, we placed sequences necessary for probe production (PCR primer amplification sites and T7 promoter), as described below. All sequences for encoding probes can be found in **Table S3**.

### Encoding probe preparation

The protocol for probe production is described on a Protocols.io webpage (https://www.protocols.io/view/highly-amplified-multiplexed-fluorescence-in-situ-ditu4enw). Of note, encoding probes were constructed with an improved protocol. Briefly, the oligo pools were amplified by limited-cycle PCR (40 ng per reaction) using KAPA HiFi polymerase (Roche, KR0369) (10 mM dNTP, 1x KAPA HiFi fidelity buffer) with the following protocol: 98**°**C for 3 min, then 22 cycles of 98**°**C for 10 s and 72**°**C for 30 s, followed by 72**°**C for 5 min. In total, four sub-libraries were amplified with the following four primer pairs; lib1_fwd (5′- GCGTTGCGTGCTAACTCGGA-3′) and lib1_rev (5′-TTACTAACCCGGTCGTGCGG-3′), lib2_fwd (5′- GAGTGCGGATACGTCGTCGT-3′) and lib2_rev (5′-AGGGACGCTTAAGTCACGGC-3′), lib3_fwd (5′- GCAGTTCGTGCGACCGTGTA-3′) and lib3_rev (5′-ATCGCACGGTTCGTAATCCG-3′), lib4_fwd (5′- TCCGTTAACGTTACGCGGTG-3′) and lib4_rev (5′-AGTGGCGCCGAATAAGCAAT-3′).

Using the HiScribe T7 High Yield RNA Synthesis Kit (New England Biolabs, E2040), the sense strand was amplified further and we then produced antisense ssDNA by reverse transcription (RT) with Maxima H Minus Reverse transcriptase (Thermo Fisher, EP0751). To increase encoding probe tissue penetration, RT primers contained RNA bases to ensure degradation of the shared RT sequences in the final encoding probes upon hydrolysis. The RT primer sequences used are as follows: lib1_RT_fwd (5′- GCGTTGrCrGTGCTAArCTCGGArU-3′), lib2_RT_fwd (5′-GAGTGCGrGrATACGrUCGTCGTrU-3′), lib3_RT_fwd (5′-GCAGTTrCrGTGCGACrCGTGTArU-3′), lib4_fwd_RT (5′- TCCGTTrArACGTTACrGCGGTGrU-3′) where rX denotes an RNA base. Our final probe length following RT sequence degradation was therefore 58-62 bases rather than 78-82. The RNA was then hydrolyzed (0.5M NaOH, 0.25M EDTA, incubated at 95**°**C for 10 min) and the reaction was neutralized with 0.33x volume of 1M Tris-HCl pH 7.0. We purified the ssDNA probes by ethanol precipitation (0.5x volume of 7.5M ammonium acetate and 2.5x volume of 100% ethanol) and resuspended the pellet in ultrapure water.

### Amplifier design and screening

Sequences used for amplifier and readout oligonucleotides were based on a collection of orthogonal 25mer DNA barcodes^42^. For the sequences used for preamplifier (19 nt) and amplifier (18 nt) repeats we shortened these sequences and selected those with a Tm range of 50-55**°**C. Fluorescent-readout probes were 25 nt, despite binding sites on amplifier oligonucleotides being only 18 nt. This allowed us to strip fluorescent-readout probes during hamFISH experiments using 25 nt ‘toehold’ oligonucleotides that are complementary to the fluorescent-readout probes. Additionally, amplifiers were synthesized with a 3′ amino modification for covalent linkage to the hydrogel. We developed 32 orthogonal sets of amplifiers over three rounds of tests. First, we designed 16 amplifier sets (consisting of preamplifier, amplifier, and fluorescent-readout probes), which we found to produce signals in varying intensities. We subsequently tested additional 18 sets to compare signals with the original 16 sets using *Sst* as the common target. Signal intensity and signal-to-noise ratio were evaluated using a Zeiss AxioImager 2 using a Plan Apo 20x objective (NA 0.8) and Orca Flash v2 (Hamamatsu Photonics) camera. In a third iteration of screening, we devised a method to screen 48 candidates in a more systematic fashion. First, we screened the bridge sequences using a fixed amplifier set. This was done by using *Sst* encoding probes with candidate bridge sequences and bridge probes that contain a common preamp binding sequence for the readout. After selecting bridge sequences that produced highly specific signals, we screened the preamplifier repeat sequences using a common *Sst* encoding probe and a bridge probe that contains a candidate preamplifier sequence. We then used another oligonucleotide that contains a sequence complementary to this candidate sequence plus 4x of the amplifier binding site. The signal was read out using a common amplifier and a fluorophore-conjugated probe. Using the screened sequences, we designed new amplifier sets consisting of preamplifier, amplifier, and label probes (label probes were not screened). Finally, we tested these new sets using *Gad1* encoding probes and corresponding bridge probes containing binding sites for 4 different amplifier sets plus an existing set as a standard. We then chose the new amplifier sets that perform comparably to the reference.

For synthesis of longer amplifiers for use in highly amplified v2 smFISH, preamplifier and amplifier sequences containing 4 amplifier-binding and readout probe-binding repeats respectively, were synthesized by overlapping PCR amplification of three primers using KAPA HiFi polymerase (Roche, KR0369) (10 mM dNTP, 1x KAPA HiFi fidelity buffer) with the following protocol: 95**°**C for 3 min, then 30 cycles of 95**°**C for 30 s and 65**°**C for 30 s, followed by 72**°**C for 3 min). The repeated sequences were also flanked with inverted nicking endonuclease (Nt.BbvCI) sites (underlined) and contained a T7 promoter sequence (italicized). Taq Polymerase (Invitrogen, K461020) was used for this, according to the manufacturer’s protocol, for its compatibility with TOPO cloning. This resulted in the following sequences: preamplifier_4x = CATTCGGTTGGGTATCGGTAATCAGCGT<u>CCTCAGC</u>(GCCTAAATTGACAGATTCC)_4_<u>GCTGAGG</u>*CC CTATAGTGAGTCGTATTA*, generated from the primers preamp_fwd1 (5′- CATTCGGTTGGGTATCGGTAATCAGCGTCCTCAGCGCCTAAATTGAC-3′), preamp_fwd2 (5′- CCTCAGCGCCTAAATTGACAGATTCCGCCTAAATTGACAGATTCCGCCTAAATTGACAGATTCCGC CTAAATTGACAGATTCCGCTGAGG-3′) and preamp_rev1 (5′- TAATACGACTCACTATAGGGCCTCAGCGGAATCTGTCAA-3′), amplifier_4x = GGAATCTGTCAATTTAGGC<u>CCTCAGC</u>(GATCTGCTATCCGTTAGT)_4_<u>GCTGAGG</u>*CCCTATAGTGAGT CGTATTA* (generated from the primers amp_fwd1 (5′- CTCTCAATGCGCAATTAAGCCTCAGCGAGCCACGTTGA-3′), amp_fwd2 (5′- CCTCAGCGAGCCACGTTGAATTATCGAGCCACGTTGAATTATCGAGCCACGTTGAATTATCGAGCCACGTTGAATTATCGCTGAGG-3′) and amp_rev1 (5′- TAATACGACTCACTATAGGGCCTCAGCGATAATTCAACGT-3′). These were cloned into pTOPO using the TOPO cloning method (Invitrogen, K461020).

Following plasmid DNA preparation and confirmation of successful clones by sequencing, the supercoiled portion of the DNA was separated from any nicked or single-stranded DNA by purification after electrophoresis on a 1% agarose gel (50 µg plasmid DNA loaded). Purified supercoiled DNA was then incubated with Nt.BbvCI (New England Biolabs, R0632) according to the manufacturer’s protocol followed by column purification. To duplicate amplifier repeats within the nicked plasmid DNA, we incubated the samples with DNA polymerase I, Large (Klenow) fragment (New England Biolabs, M0210) which fills in 5′ overhangs (left by the nicking endonuclease) in the presence of single-stranded binding proteins (SSBP) to stabilize secondary structure (2 µg DNA, 1x NEBuffer 2, 1.6 mM dNTPs, 5 µg SSBP, 2.5 units Klenow, 37°C for 1h). The resulting single-stranded (filled-in) DNA was then gel purified and ligated with T4 DNA ligase (New England Biolabs, M0202) before transformation into Stbl3 competent cells (Thermo Fisher, C737303). Plasmid DNA that had been successfully nicked and filled-in therefore had amplifier 8 repeats, which was confirmed by sequencing. This process was repeated 2 more times, resulting in plasmid DNA with 32 repeats of either amplifier-binding or readout probe-binding sequences.

To synthesize single stranded amplifier sequences from these plasmids, DNA was digested with EcoRI to separate the amplifier template from the plasmid backbone. This template was gel purified and used as a template for *in vitro* transcription (New England Biolabs, E2040). The resulting RNA was reverse transcribed with Maxima H Minus Reverse transcriptase (Thermo Fisher, EP0751). Note that for the amplifier sample, dNTPs were supplemented with 100 nM aminoallyl-dUTP (Thermo Fisher, R0091) for incorporation into the tissue hydrogel during gel embedding. The RNA template was removed by alkaline hydrolysis, as described in the probe preparation methods above, and the remaining ssDNA was neutralized with 0.33x 1M Tris-HCl pH7.0, then column purified (Omega Biotek, D6943).

### Tissue preparation for hamFISH experiments

Tissue was harvested by freezing freshly dissected brains of animals in OCT compound (Tissue-Tek, 4583). For animals injected with cholera toxin subunit b, the lobe containing the left MeA was cut away from the rest of the brain (which was drop-fixed in 4% v/v paraformaldehyde (Thermo Fisher, 28908) in 1x phosphate-buffered saline (PBS) (Severn Biotech, 20-74-10) and used for injection site confirmation, see below) before freezing in OCT. Embedded brains were stored at -80**°**C. Prior to sectioning, 40 mm diameter #1.5 coverslips (Bioptechs, 0420-0323-2) were treated with 0.5% v/v (3-mercaptopropyl)trimethoxysilane (Sigma, 175617) and 0.34% glacial acetic acid in ethanol, and poly-L-lysine (Sigma, P8920) for silanization and promotion of tissue-adhesion respectively.

Frozen brains were manually sectioned onto these coverslips at -18**°**C on a cryostat (Leica, CM3050S). Surrounding tissue was trimmed to ensure approximately 18-22 sections (9-11 from each animal, 2 animals per coverslip) could fit onto the imaging area of each coverslip. Sections were cut at 10 µm thick with every 10^th^ section mounted, ensuring representative sections from the full anterior-posterior axis of the MeA was present (Bregma - 1.2 mm to -2.2 mm).

Samples were then fixed with 4% v/v paraformaldehyde, 0.4% v/v glyoxal (Sigma, 50649), 0.1% v/v methanol in 1xPBS^43^ followed by treatment in pre-chilled 100% ethanol at -80**°**C for 30 min. Subsequent drying out of these samples was followed by rehydration in 2x saline-sodium citrate (SSC) (Severn Biotech, 20-6500-50) and clearing in 4% wt/v sodium dodecyl sulfate (SDS) (Sigma, 75746) in 1x PBS for 5 min each, on a rotating platform. After three quick washes in 2xSSC, sections were washed with 1x TE (Severn Biotech, 20-5310-05) (2 washes at 1 min each) followed by permeabilization using protease (0.34 µg/ml) (Sigma, P5380) in 1x PBS for 7.5 min. Samples were then washed again in 2x SSC followed by a final wash in 2x SSC containing 20% v/v formamide (Sigma, F7503).

### Encoding probe staining, amplification, and gel embedding

The molarity per gene was estimated based on the number of genes in the probe library, despite some differences in the number of probes per gene. For example, a probe library containing 30 genes with a total concentration of 30 µM ssDNA was defined as 1 µM per gene and diluted accordingly. Encoding probes were diluted in hybridization buffer; 2x SSC, 22.5% v/v formamide (Sigma, 47671), 5% wt/v dextran sulfate (Millipore, S4031), 0.5% v/v TWEEN-20 (VWR, 437082Q), 10 mM ribonucleoside vanadyl complexes (New England Biolabs, S1402), 0.1% wt/v yeast RNA (ribonucleic acid from Torula utilis, Sigma, 83850). hamFISH encoding probe libraries were used at a final concentration of 25 nM per gene as defined above, and individually synthesized probes at 100 nM per gene. A polyA-anchor probe containing locked nucleic acids (LNA) as described in ref^23^ was also included at a concentration of 1 µM (/5Acryd/TTGAGTGGATGGAGTGTAATT+TT+TT+TT+TT+TT+TT+TT+TT+TT+T). This probe permits RNA anchoring to a polyacrylamide gel as described below.

Coverslips were inverted onto a drop of hybridization buffer (approximately 160 µl) and placed onto a layer of parafilm sitting in a 10 mm petri dish, with a 0.75 mm thick spacer to ensure even hybridization. The sample was incubated at 40**°**C for 16-18 h before being washed in 2x SSC, 20% v/v formamide (wash buffer), initially at room temperature for 3 quick washes, followed by incubation for 1h at 40**°**C. For signal amplification samples were incubated in sequential hybridization of bridge-readout, preamplifier, and then amplifier oligonucleotides in amplifier hybridization buffer (2x SSC, 20% v/v formamide, 10% wt/v dextran sulfate) for 30 min at 40**°**C. This was interspersed with 2x 5 min washes in wash buffer on a rotating platform. Bridge-readout and blocking probes were used at a concentration of 1 nM per gene, whereas preamplifiers and amplifiers were used at 10 nM per set. After the amplification and wash steps, samples were fixed again for 10 min with 4% v/v paraformaldehyde in 1x PBS followed by a 2x SSC wash, then either frozen at -80**°**C after drying or processed for gel embedding and signal development (see below).

For gel embedding, samples were treated as previously described^11^. Briefly, coverslips were washed in 20 mM acrylic acid NHS ester (Sigma, A8060) for 90 min, followed by a 30 min incubation in monomer buffer; 2x SSC, 4% acrylamide and 0.2% bis-acrylamide (Bio-Rad Laboratories). Aspirated samples were then inverted onto a 40 µl drop of polymerization mixture (monomer buffer with the addition of 0.2% ammonium persulfate and 0.2% tetramethylethylenediamine) upon a Gel Slick (Lonza, 50640) treated glass plate. After polymerization, coverslips were removed from the glass plate while submerged in 2x SSC, and we proceeded with digestion with 1 mg/ml proteinase K (Ambion, AM2548) in 2x SSC, 2% wt/v SDS, 0.5% triton-X (Sigma, X100) for 3 h at 40**°**C on a rotating platform.

### Imaging platform

Samples were imaged on a custom-built imaging platform based on that described by^22, 44^ with 405, 488, 568 and 647 nm solid-state lasers (OBIS, Coherent). The base of the microscope was Nikon Ti-U (Nikon) equipped with a custom FocusLock system as described in^22^. The coverslip was assembled in a flow chamber (Bioptechs, FCS2) attached to a home-built fluidics system that controlled hybridization flow through the sample. The fluidics system comprised of a peristaltic pump (Gilson, MINIPULS 3) and three eight-way valves (Hamilton, MVP RS-232 valves), all computer-controlled by HAL/Dave and Kilroy software (https://github.com/ZhuangLab/storm-control/blob/master/storm_control/README.md) ensuring fully automated control of the whole system.

### Imaging

The flow chamber was filled with imaging buffer during acquisition (2x SSC, 50 mM Tris-HCl pH8.0, 10% wt/v glucose (Sigma, G8270), 2 mM 6-Hydroxy-2,5,7,8-tetramethylchromane-2-carboxylic acid (Sigma, 238813), 250 µg/ml glucose oxidase (Sigma, 345386), 40 µg/ml catalase (Sigma, C30) and 40 U/ml RNase inhibitor (New England Biolabs, M0314)). Each hybridization readout buffer (2xSSC, 40% v/v formamide, 5% wt/v dextran sulfate) contained AlexaFluor-488, Cy3 or Cy5 fluorophore-conjugated readout probes (50 nM each) and toehold probes (1 µM each) corresponding to the previous imaging round’s fluorescent readout probes. The hybridization protocol for each round was as follows (at an approximate flow rate of 0.15 ml/min); 600s readout and toehold hybridization solution, 1800s stop (incubation time for hybridization), 350s 2x SSC, 100s stop, 180s 2x SSC with 8 ng/µl DAPI, 180s 2x SSC, 150s stop, 200s imaging buffer. Note that a DAPI step was included in our protocol to supplement DAPI signals that became weaker in each round. For barcoded hamFISH, a 60x Plan Apo 1.4NA oil objective (Nikon) was used for imaging. A Z-stack for each channel was acquired at each field of view, covering a thickness of 7 µm with 14 steps at 500 nm each. Typically, 30-40 FOVs covering 1.25-1.65mm^2^ were needed to fully image each MeA section. The imaging area for each coverslip (containing the MeA from 1 animal, 12 sections from each) was therefore approximately 17mm^2^. A 40x Plan Apo 1.3NA oil objective (Nikon) was used for imaging. A Z-stack for each channel was acquired at each field of view, covering a thickness of 5 µm with 6 steps at 1 µm each. Typically, 12 (4x3) FOVs covering 1.25mm^2^ were needed to fully image each MeA section. The imaging area for each coverslip (containing the MeA from 2 animals, 10 sections from each) was therefore approximately 25mm^2^.

### Stereotaxic injection of retrograde tracers

Cholera toxin subunit b (CTB) -AlexaFluor488 (Thermo Fisher, C34775), CTB-AlexaFluor555 (C34776), and CTB-AlexaFluor647 (C34778) were used for retrograde tracing of MeA-projection neurons. Briefly, adult male CD1 mice were anesthetized in an induction chamber containing 4% isoflurane mixed with oxygen. After shaving the head, animals were then transferred to a stereotaxic frame and anesthesia was maintained with ∼1% isoflurane delivered through a nose cone. A small incision was made and a hole was drilled at each of the injection sites. CTB (0.5%, 75nl) was delivered through pulled glass micropipettes to the target location using Nanoject III (Drummond Scientific); MPOA (AP -0.1 mm, ML -0.5 mm, DV +5.4 mm), BNST (AP -0.2 mm, ML -1.1 mm, DV +4.3 mm), VMHvl (AP -1.4 mm, ML -0.7 mm, DV +5.9 mm), PMv (AP -2.5 mm, ML -0.5 mm, DV +5.8 mm). After injections, the scalp was sutured and animals were allowed to recover in their cage placed on a heated platform for 1h. Mice were euthanized by cervical dislocation 5-7 days after CTB injection. After dissection of the lobe containing the left MeA (which was frozen in OCT compound), the rest of the brain was drop-fixed in 4% v/v paraformaldehyde 1x PBS overnight at 4**°**C, before vibratome sectioning at 75 µm thickness and mounting with DAPI-containing mounting medium: 80 ng/µl DAPI (Sigma, D9542), 0.2% v/v n-propyl gallate, 90% v/v glycerol, 1x PBS. Injection sites were confirmed using Zeiss Axio Scan imaging.

### Prehybridization imaging of CTB signal in the MeA

The left MeA lobe (previously frozen in OCT, to be used for FISH) was sectioned as described above. After fixation in 4% v/v paraformaldehyde, 0.4% v/v glyoxal, 0.1% v/v methanol in 1xPBS, and staining with 2x SSC 80 ng/µl DAPI, samples were washed in 2x SSC containing RNase inhibitor (40 U/ml, New England Biolabs, M0314) then mounted onto slides. Sections were then imaged using a Zeiss Axio Imager 2 using the 20x objective and Orca Flash v2 (Hamamatsu Photonics) camera, with a 4-step Z-stack covering 12 µm thickness. After imaging, slides were submerged into 2x SSC to loosen the coverslip without damaging the sample, and the tissue preparation protocol described above was followed starting with the ethanol incubation step.

### Transcriptomic analysis pipeline

To analyze the previously published scRNA-Seq data^8^, clustering was performed using Seurat^45^. We set the number of dimensions to 10 and a resolution of 2. After quality control (set as requiring 200-2500 RNA per cell), we selected the top 250 genes that showed significant variation for clustering.

For hamFISH experimental data, we built a smFISH analysis pipeline that creates maximum intensity projections from each imaging round (ImageJ). To correct the illumination bias of the excitation beams in the field of views, the images were corrected using a blurred average image compiled from all tiles across all hybridization rounds for each channel to calculate intensity differences. These images were then used to normalize intensity differences for DAPI images. This was then combined with DAPI-based Cellpose 2.0 cell segmentation^46^. To account for slight shifts in the stage position across hybridization rounds, DAPI signal was used to calculate the translation in XY for every tile. This was done using the phase cross-correlation of *scikit-image*^10^ (https://github.com/khchenLab/split-fish). The modal value for each hybridization round was then taken and applied to all images for that round, as well as to Cellpose masks to define individual cell positions across the experiment.

A Fast Fourier transform (FFT) based frequency filter (200Hz) was applied to signal images (i.e., non - DAPI channels) to exclude background haze. These then underwent shading correction as described above for DAPI. The intensity of signal within each cell (chosen by a manually defined intensity cut-off) was then used to define positive and negative cells for each transcript. The intensity value below the threshold was subtracted, and the intensity was divided by the standard deviation of each bit to adjust for the differences in variance in each bit. This data is therefore comparable to normalized cell x gene data. To test for orthogonality, the centroids of each transcript signal were found using 3D-DAOSTORM^47^ and overlap was determined to be positive if two or more bits were within 0.25 pixels. For CTB-injected smFISH experiments, the DAPI signal from hamFISH experiments was aligned to the prehybridization CTB-imaging DAPI signal using a custom Python script that performed affine transformation to account for distortion (cv2.warpAffine). We then manually verified the alignment of each section. All code for hamFISH data processing is available at https://github.com/mathew-d-edwards/hamFISH.

Clusters were determined using a combination of iterative Seurat-based clustering and manual confirmation with hamFISH signal overlays. Briefly, Seurat was used to define an initial set of clusters which were then checked by manual overlays of signal from differentiating genes confirm clusters. After further manual overlays with other genes, higher-resolution cluster definitions could be determined. After cells that fit into these determined clusters were removed from the analysis, the remaining cells were subject to further Seurat-based clustering to try and determine their type. This iterative process gave an initial list of clusters that we then tried to find in multiple datasets and allowed us to ensure clusters were consistent in number across animals and experiments. A final cluster definition was then used to assign all cells from every experiment into clusters. The data was then analyzed and presented using R, Sklearn (t-SNE), Scanpy (violin plots)^48^, Seurat (heatmaps), GraphPad Prism, and Python-based plotting scripts for cluster spatial plots. Comparison of scRNA-Seq clusters with hamFISH clusters was done using Pearson correlation between group averages of the expression of each transcript in our gene marker panel.

For spatial analysis of hamFISH clusters, we defined several parameters that were used to cluster cell types based on their spatial features. To do this we identified common sections across samples, first in z by manually defining anterior-posterior Bregma position based on cluster locations. Then in x and y by manually defining the ventral and medial edge of the MeA, as well as the most ventral-medial point that the MeA reaches, and by using the angle between all three to align sections between samples. Spatial features used to cluster cell types were: mean x (medial-lateral axis) position, standard deviation (SD) of x, mean y (dorsal-ventral axis), SD of y, mean z (anterior-posterior axis), SD of z, total number of cells, distance to molecular layer (manually defined in reference images), proportion of cells with nearest neighbor cells belonging to cluster i1, and proportion of cells with nearest neighbor cells belonging to cluster i2 etc. for all clusters. For visualization, t-SNEs (Sklearn) were generated with a perplexity of 50, n_components=2, and an automatic learning rate.

### Input/output tracing of Ndnf and Sst neurons

Following craniotomies as described above, the following AAV or rabies viruses were used for injection into the MeA (AP -1.6 mm, ML +/- 2 mm, DV +5.6 mm) of 2-3 month old NDNF-Cre or SOM-Cre male mice (C57BL/6J mice backcrossed to CD1 background). For monosynaptic retrograde rabies tracing experiments, 20-60 nl of AAV2/1.syn.flex.H2BG.N2CG and AAV2/1.flex.GFP-TVA (6.94E+13 vg/ml and 3.84E+13 vg/ml, respectively; Francis Crick Institute viral vector core) were injected unilaterally into the MeA followed two weeks later by 20-60nl of N2C△GEnvA.Rabies.mCherry (1E+09 vg/ml; Sainsbury Wellcome Centre viral vector core (SWC-VVC)). To determine the projection targets of Ndnf cells, 100nl of AAV2/2.EF1a.FLEX.Synaptophysin-EGFP and AAV2/2.CAG.FLEX.tdTomato mix (6.93E+13 vg/ml and 5.77E+13 vg/ml, respectively (SWC-VCC)) were injected into the MeA of 8-12-week-old male CD1 mice. Tissue was harvested 4-5 weeks after injection. After perfusion, dissection, and post-fixation of the brain in 4% v/v paraformaldehyde 1x PBS overnight at 4°C, brains were embedded in agar and imaged using serial section two-photon microscopy^49, 50^. Imaging was controlled by ScanImage Basic (Vidrio Technologies, USA) with BakingTray, a custom software wrapper for setting up imaging parameters (https://github.com/SainsburyWellcomeCentre/BakingTray). Sections were cut at 50 µm and the signal was imaged with 780 nm excitation wavelength. Subsequent image assembly was performed using StitchIt (https://github.com/SainsburyWellcomeCentre/StitchIt). Sections were then registered to the 10 µm Allen Brain Atlas using BrainReg software^51, 52^. For the analysis of our input mapping, cellfinder software was used to automatically detect input cells^53^. Although a low error rate of detection was seen for most areas, the main and accessory olfactory bulbs consistently gave an unacceptable error rate due to high background levels and inaccurate registration and therefore input cells were manually counted in these areas. For the analysis of our output mapping, the puncta-like signal from Synaptophysin-EGFP did not allow for automatic cell detection with cellfinder. The serial section two-photon microscopy images were therefore used to identify regions of interest, before re-mounting and imaging these areas using a Zeiss Axio Scan. These images were registered to the Allen CCFv3 atlas using AP_histology software (https://github.com/petersaj/AP_histology) and the signal was then manually counted as puncta using ImageJ to give a density value for each area of interest.

### Hybridization Chain Reaction (HCR)

All HCR sequences can be found in Table S4. Briefly, Ndnf, Sst and Chrna7 encoding probes were designed with overhangs complementary to bridge sequences that served as initiators for hairpin amplification. HCR was performed using the same tissue preparation protocol used for hamFISH (see above) except that 20 µm thick sections were used instead of 10 µm, and samples were fixed for 10 min at room temperature in 1x PBS with 4% v/v paraformaldehyde. After tissue preparation, the manufacturer’s protocol (Molecular Instruments) was used to develop the HCR signal with the following alterations. We used encoding probes at a final concentration of 10 nM for our initial overnight incubation. On the following day a bridge probe hybridization step was performed in HCR Hybridization buffer for 30 min at 37**°**C with a bridge probe concentration of 10 nM. The section was washed with 0.2x SSC to remove the unbound bridge probe. To develop HCR signal, after denaturation (95**°**C for 90 sec) and cooling, B2-549 and B4-647 (30 nM each) were hybridized in an Amplification buffer (Molecular Instruments) overnight at room temperature before final washes, mounting and imaging.

### Rolling Circle Amplification (RCA)

All RCA encoding probe sequences can be found in Table S4. Samples were prepared and processed as previously described^11^. A final encoding probe concentration of 100 nM was used and incubated overnight at 40°C. Probes included the complementary sequence for bit 25 fluorescent-readout probe so signal could be developed.

## Supporting information

Amplifier sequences

scRNASeq clusters

Sequential hamFISH probe sequences

HCR sequences

## Acknowledgements

We thank the Sainsbury Wellcome Centre Neurobiological Research Facility for technical support; the Sainsbury Wellcome Centre Advanced Microscopy Facility for help with serial 2-photon microscopy; and Johannes Kohl for manuscript feedback. This work was supported by the Sainsbury Wellcome Centre core grant from the Gatsby Charitable Foundation and Wellcome Trust (090843/F/09/Z).

## Author contributions

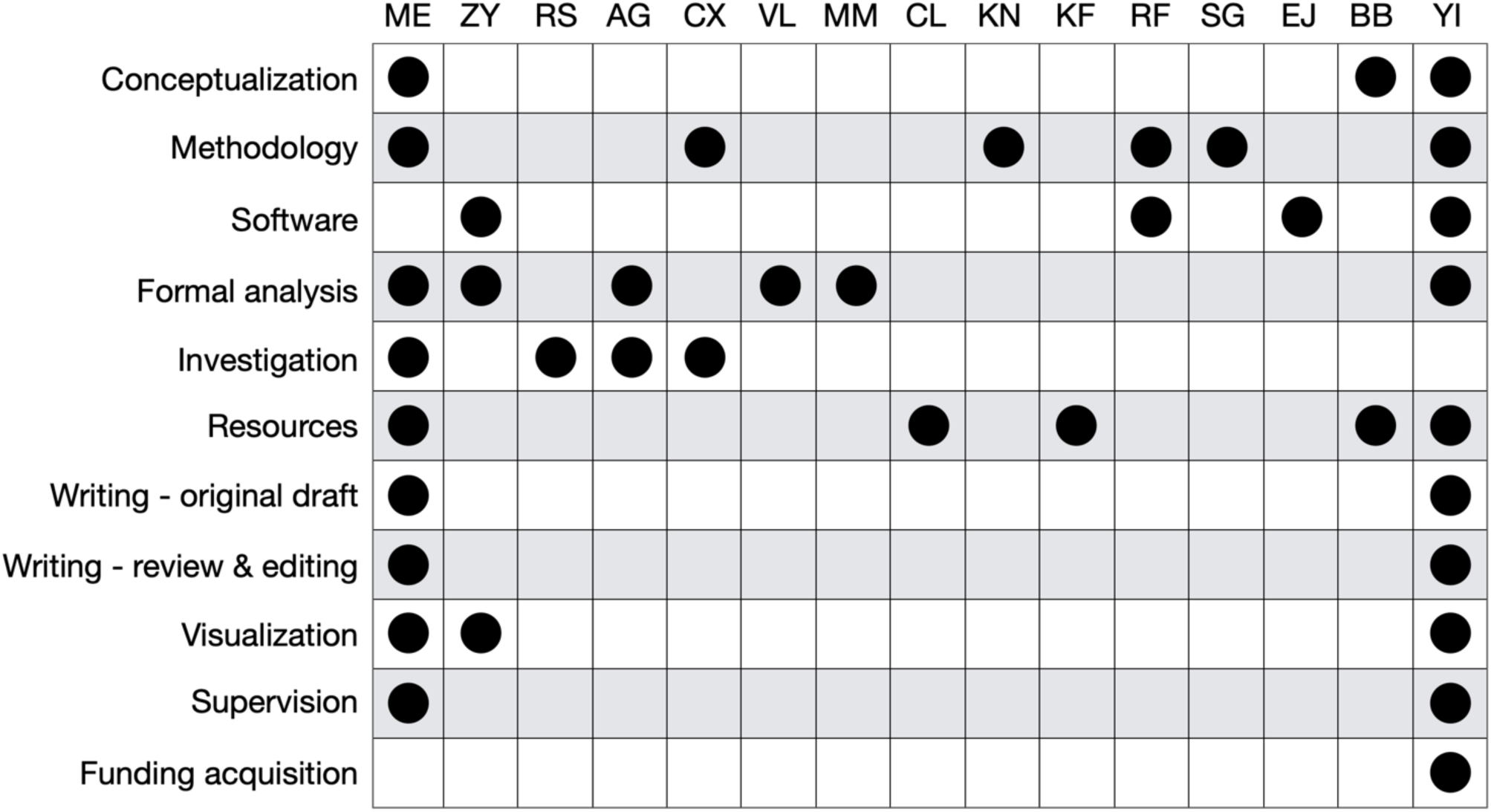

## Supplemental figures

**Supplemental Figure 1.**
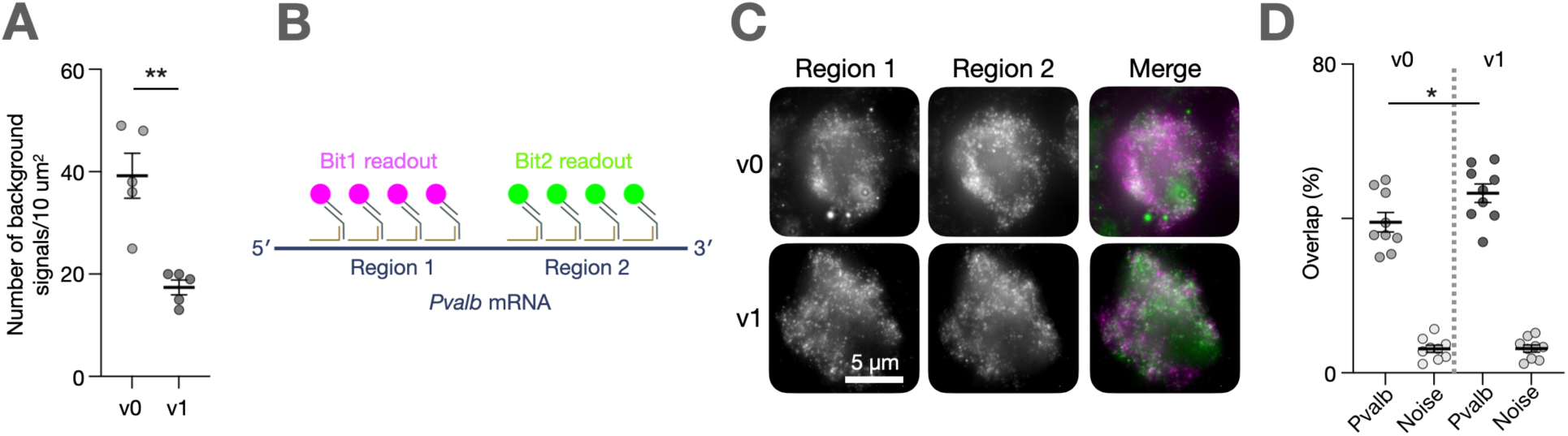
Amplification scheme v1 reduces background signals and increases hybridisation efficiency versus amplification scheme v0. **A.** Comparison of background signal abundance between v0 and v1 amplification schemes. n = 5 background areas (each 10 um2), p < 0.01, unpaired T-test. Note that left lobe (containing the MeA) was dissected for cell type profiling so is not present in these sections. Green = CTB-Alexafluor 488 conjugate, Red = CTB-Alexafluor 555 conjugate, Blue = CTB-Alexafluor 647 conjugate, Grey = DAPI. **B.** Probes were designed to target two different regions of Pvalb mRNA (region 1 and region 2) which were each developed with a different fluorophore. **C.** Representative examples of Pvalb smFISH signal from region 1 and region 2 using the v0 and v1 probe designs. **D.** Quantification of overlap of signal from each probe design. Each data points represents the overlap of real Pvalb+ signal vs background noise. n = 3 cells/background areas per sample, n = 3 samples. Comparison of signal overlap between v0 and v1, p = 0.047, unpaired T-test.

**Supplemental Figure 1 – figure supplement 2.**
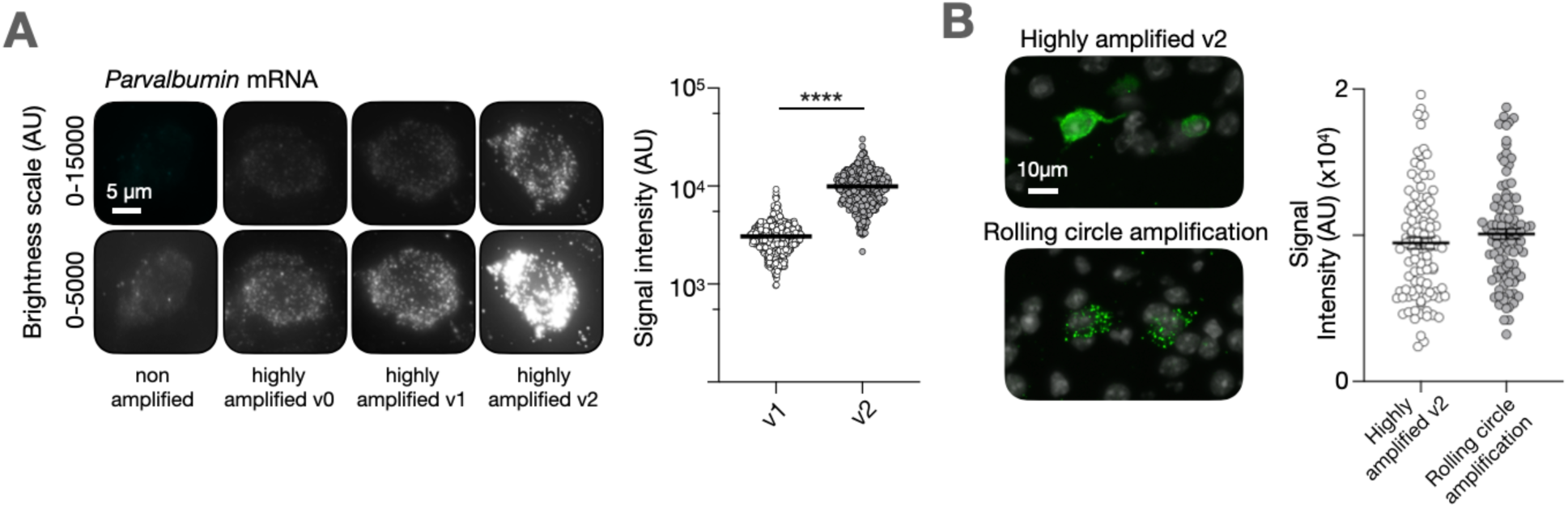
Highly amplified smFISH v2 further increases signal, making it comparable to other highly amplified methods of transcript detection. **A.** Left; representative images of Pvalb smFISH signal developed using our different amplification methods. Right; quantification of signal intensity comparison between v1 and v2, p < 0.0001, unpaired T-test. **B.** Left; representative images of Somatostatin (Sst) smFISH using with probes that were compatible with either highly amplified v2 smFISH or rolling circle amplification. Right; quantification of fluorescence intensity from each method. n = 100 each, comprising 10 ROIs from 10 cells. p = 0.23, unpaired T-test.

**Supplemental Figure 1 – figure supplement 3.**
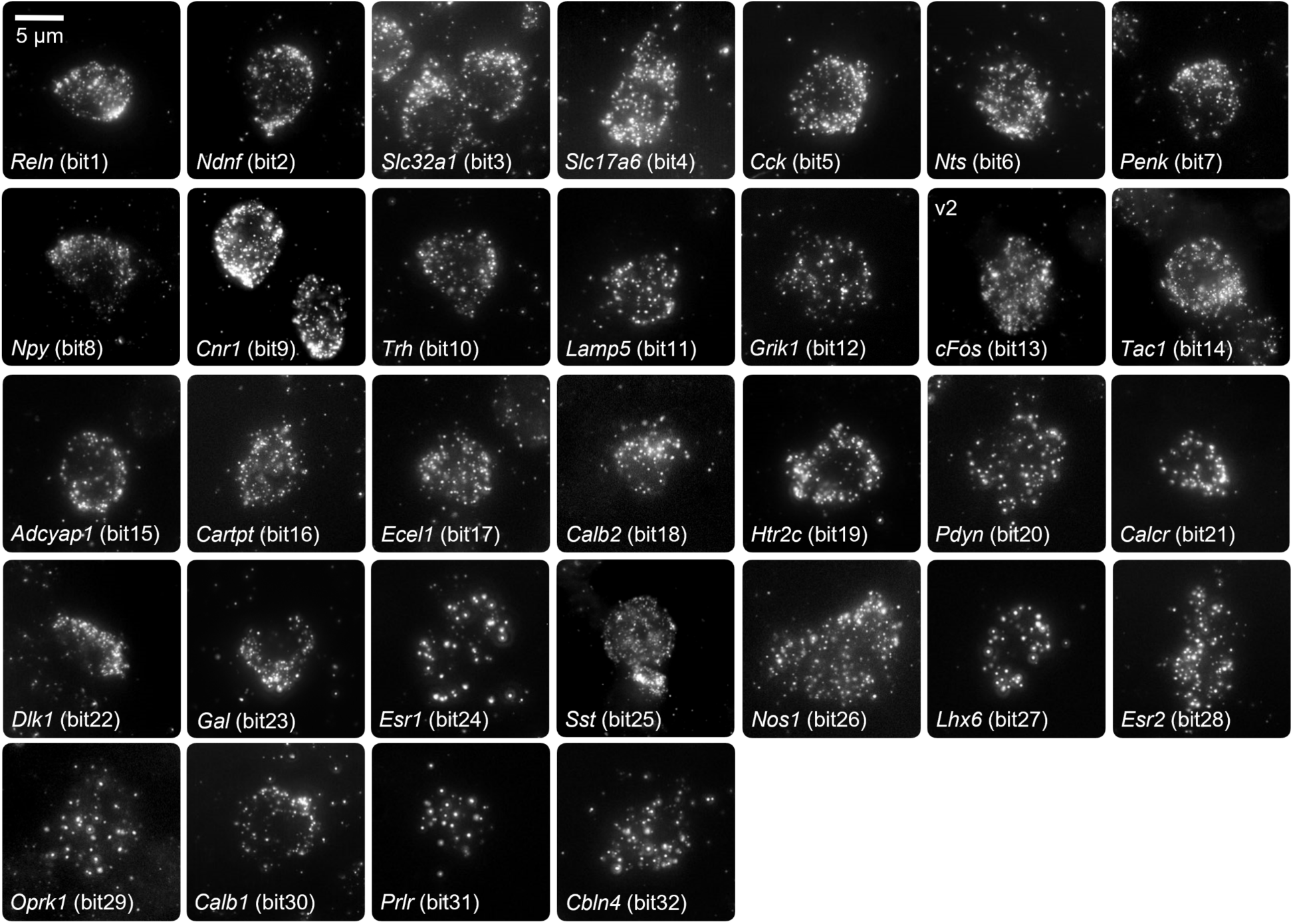
hamFISH signal using all 32 orthogonal amplifier sets. Each panel shows clear detection of individual genes by hamFISH. Each bit was used to detect a different gene. All images are taken from MeA tissue.

**Supplemental Figure 2 – figure supplement 1.**
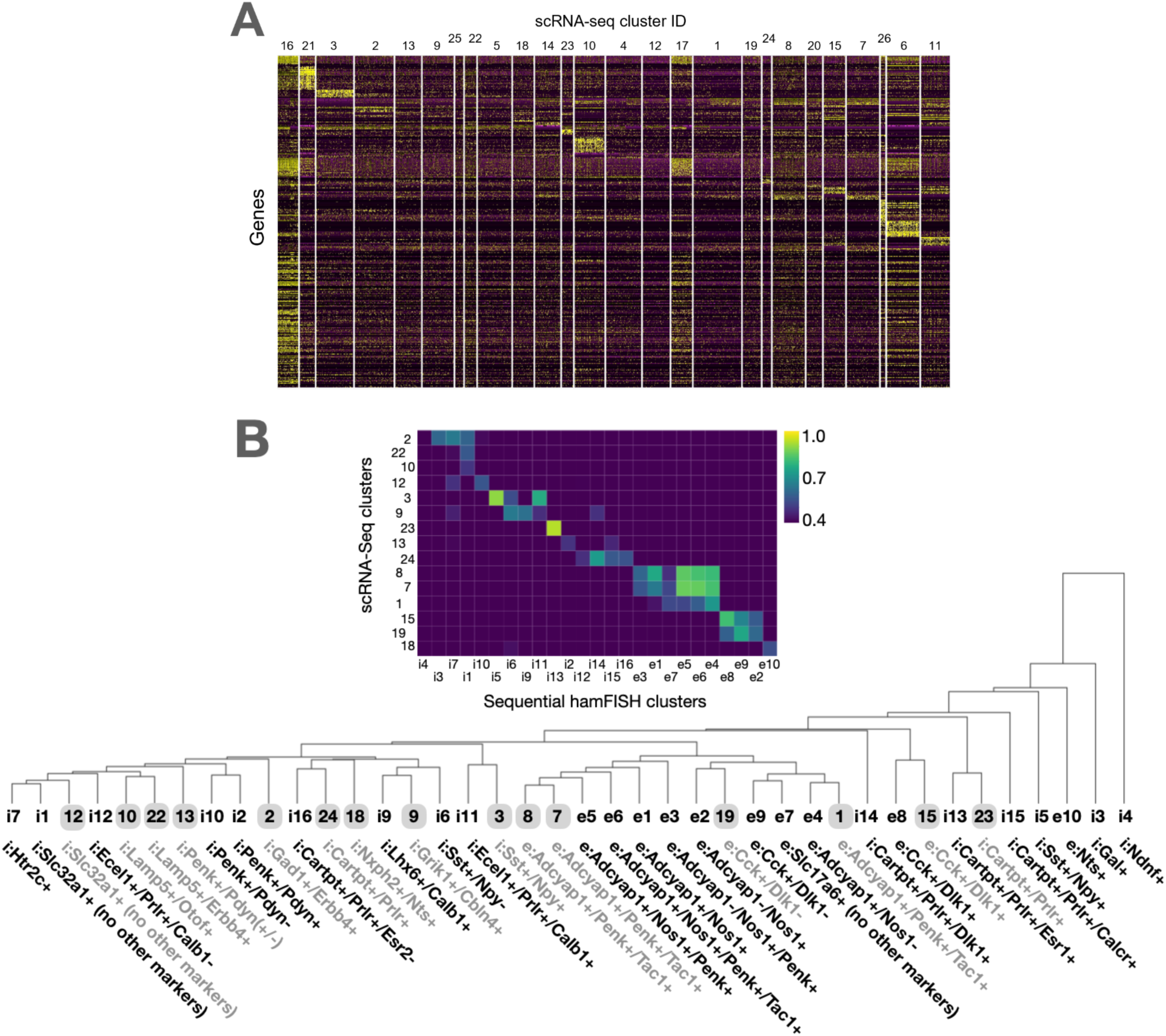
MeA clusters map onto previously published scRNA-seq data. **A.** Heatmap of clustered scRNA-seq data from Chen *et al*. **B.** Top: Heatmap showing the correlation in gene expression between scRNA-seq and sequential hamFISH clusters. Values <0.4 were excluded from analysis. Bottom: Hierarchical clustering comparing average gene expression in scRNA-seq (grey) and sequential hamFISH (black) datasets.

**Supplemental Figure 2 – figure supplement 2.**
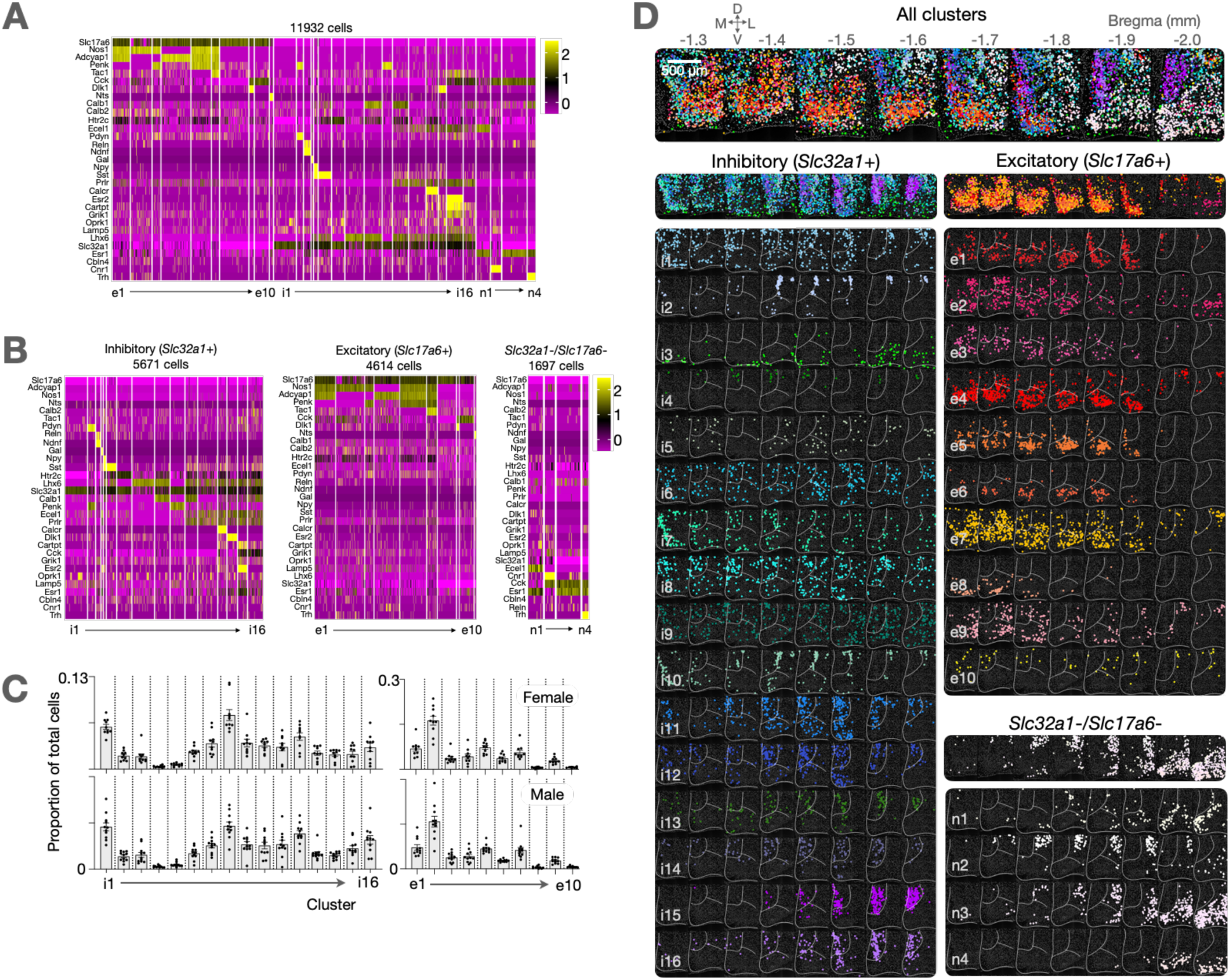
MeA cell types are consistent between animals and sexes. **A.** A combined heatmap of those shown in Figure 2A to show gene consistency across rows. **B.** Heatmap representation of the 30 identified clusters from one animal (female). Note gene order differences between plots to aid with visualisation of clusters. **C.** Comparison of number of cells per cluster as a proportion of total MeA cells. There is no significant difference between each sex. Female n = 10, Male n = 11. **D.** The spatial position of every cell from each cluster from the animal shown in B.

**Supplemental Figure 2 – figure supplement 3.**
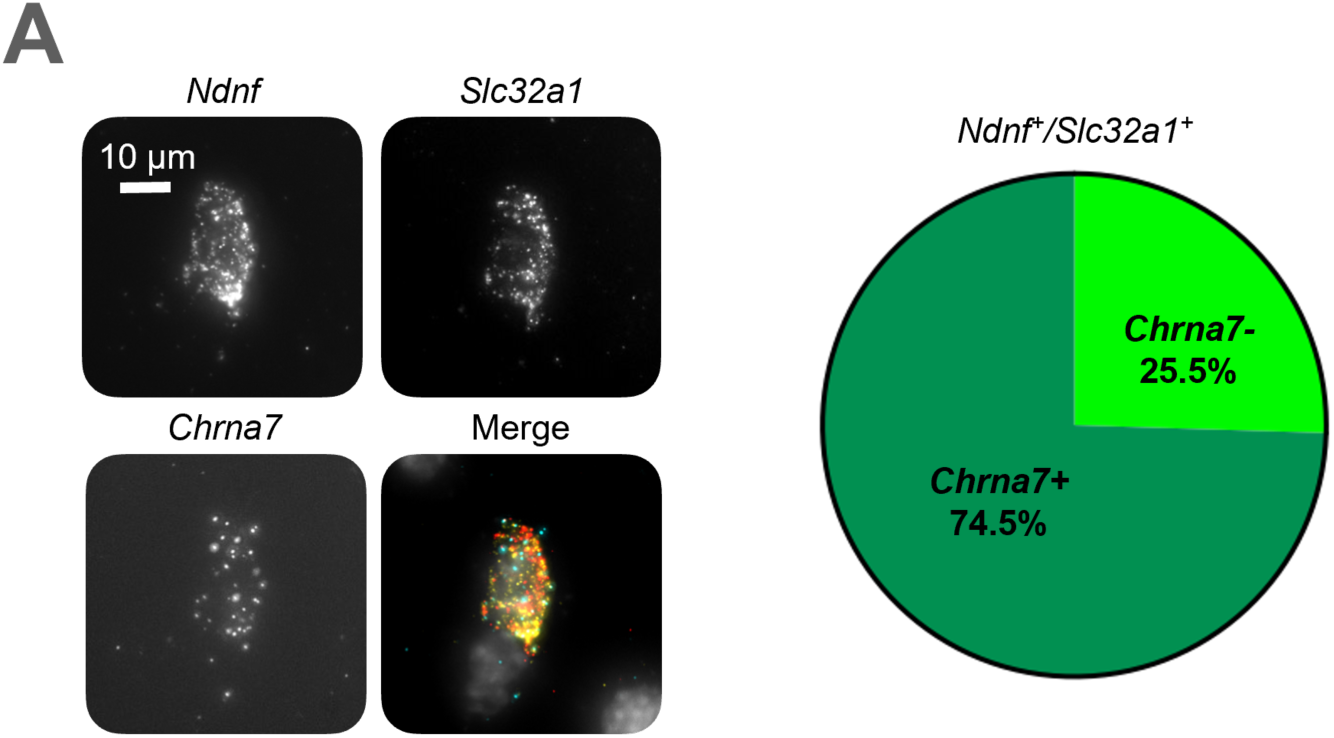
Characterization of cluster i3. **A.** A representative i3 cell with smFISH signal from *Ndnf, Slc32a1* and *Chrna7*. **B.** Of 55 *Ndnf^+^/Slc32a1^+^* neurons from 3 animals, 41 were *Chrna7^+^*.

**Supplemental Figure 2 – figure supplement 4.**
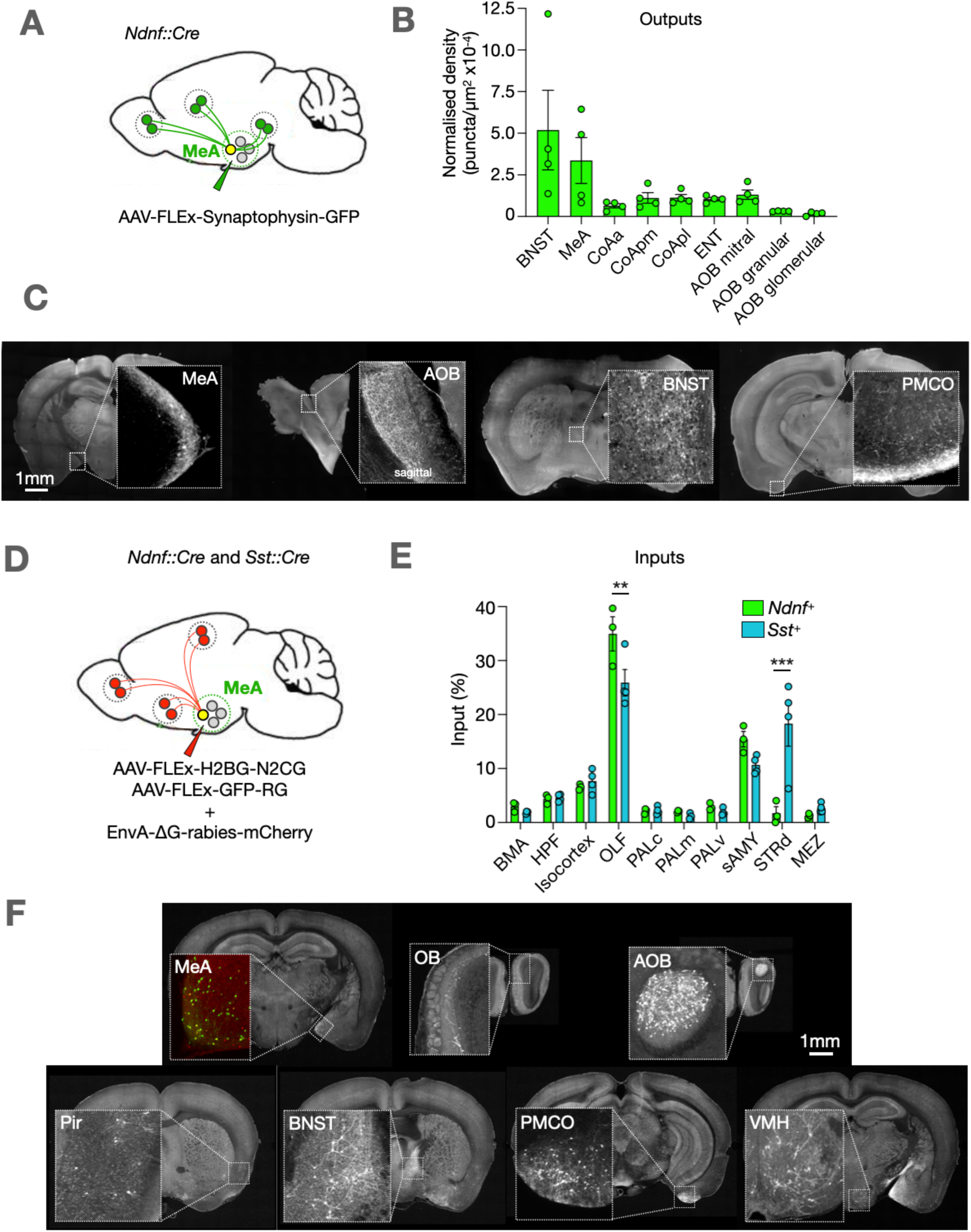
The input-output logic of MeA*^Ndnf+^* neurons. **A.** Visualization of *MeA^Ndnf+^* outputs. **B.** Projection targets of *MeA^Ndnf+^* neurons (n=4 animals). Normalized to number of starting cells per animal (n=4 animals). BNST = bed nuclei of the stria terminalis, COAa = cortical amygdalar nucleus, anterior part, COApl = cortical amygdalar nucleus, posterior part, lateral zone, COApm = cortical amygdalar nucleus, posterior part, medial zone, ENT = Entorhinal cortex, AOB = accessory olfactory bulb. **C.** Representative images of output areas with Synaptophysin^+^ neurons. **D.** Visualization of *MeA^Ndnf+^* inputs. **E.** Inputs to *MeA^Ndnf+^* (n=3 animals) and *MeA^Sst+^* (n=4 animals) neurons. Significance from 2-way ANOVA comparing Ndnf to Sst neurons. BMA = basomedial amygdalar nucleus, HPF = hippocampal formation, OLF = olfactory areas, PALc = central pallidum, PALm = medial pallidum, PALv = ventral pallidum, sAMY = Striatum-like amygdala nuclei, STRd = dorsal striatum, MEZ = hypothalamic medial zone. **F.** Representative images of output areas with rabies^+^ neurons.

**Supplemental Figure 3 – figure supplement 1.**
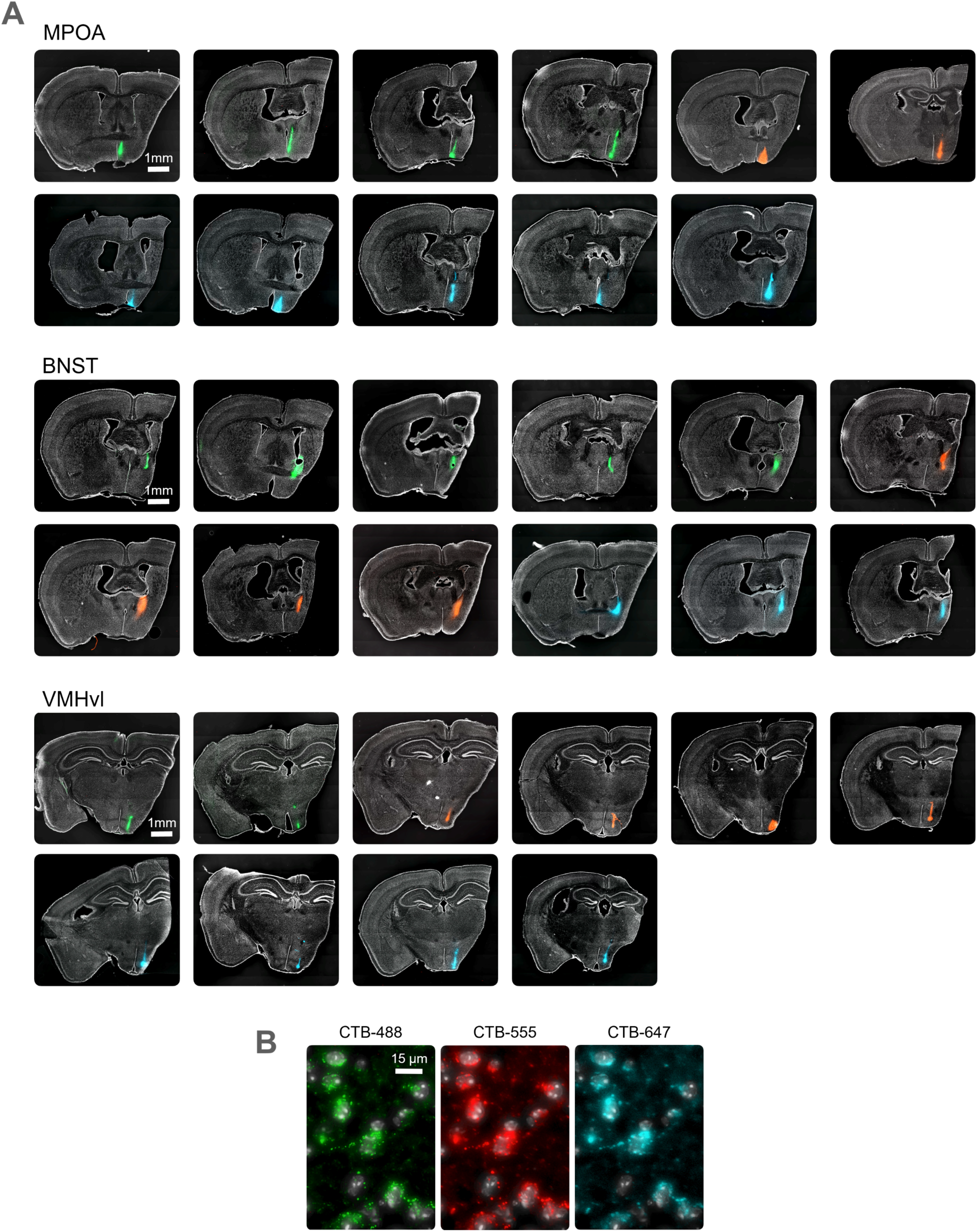
Projection cells in the MeA were labelled with CTB injections in downstream areas. **A.** Cholera toxin subunit b injection sites. Note that left lobe (containing the MeA) was dissected for cell type profiling so is not present in these sections. Green = CTB-Alexafluor 488 conjugate, Red = CTB-Alexafluor 555 conjugate, Blue = CTB-Alexafluor 647 conjugate, Grey = DAPI. **B.** MeA neurons from triple-injected BNST animal.

**Supplemental Figure 3 – figure supplement 2.**
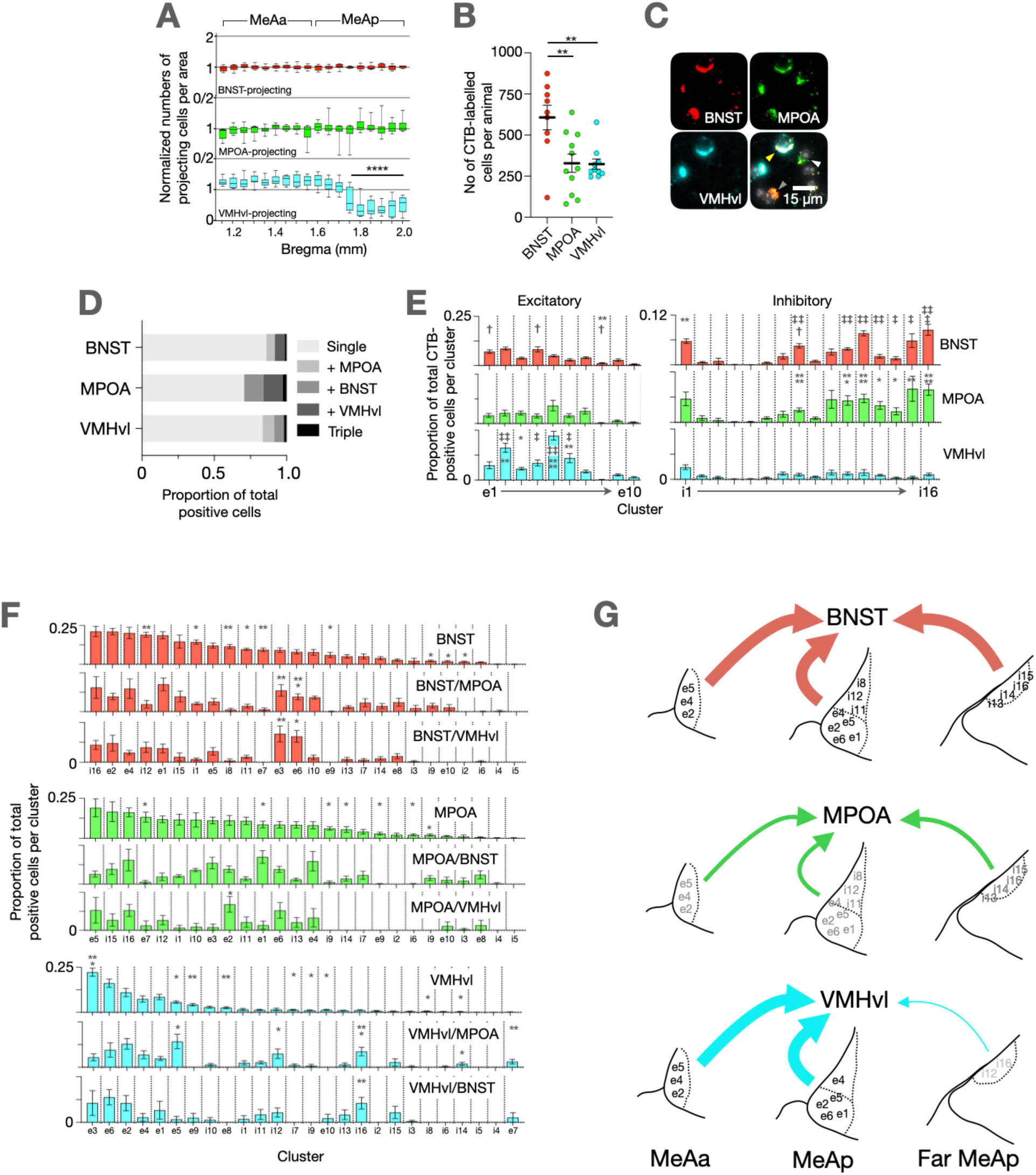
MeA projection neurons are made up of specific neuronal classes. **A.** Quantification of projecting cells along the anterior-posterior axis. For each target region, the number of CTB-positive cells at each bregma position was normalized to the total number of CTB-positive cells in that animal, with 1 being the mean. n = 8, 11, 10 for BNST, MPOA, VMHvl respectively. For VMHvl, p < 0.0001, F=26.03, one way ANOVA with a significant decrease at bregma -1.7mm (p < 0.0001, Tukey post-hoc pairwise comparison). For BNST and MPOA p=0.97, F=0.43 and p=0.20, F=1.30 respectively, one way ANOVA. **B.** Quantification of the total number of CTB-labelled neurons. p < 0.01 for BNST vs both MPOA and VMH, unpaired T-test. BNST = 621 ± 83 cells, MPOA = 341 ± 55 cells, VMHvl = 329 ± 55 cells. **C.** Representative images of single (white arrow), double (grey arrow) and triple (yellow arrow) CTB-labelled MeA neurons. **D.** Quantification of single/double/triple CTB-labelled cells. Of BNST-projecting neurons, 86±6% were single-labelled, 6±3% co-labelled with MPOA-injected CTB, 7±5% co-labelled with VMHvl-injected CTB, of MPOA-projecting neurons, 71±12% were single-labelled, 13±9% co-labelled with BNST-injected CTB, 13±11% co-labelled with VMHvl-injected CTB, of VMHvl-projecting neurons, 83±4% were single-labelled, 8±6% co-labelled with MPOA-injected CTB, 7±3% co-labelled with BNST injected CTB. **E.** CTB-positive cells per cluster as a proportion of total CTB-positive cells. Enrichment indicated by † (BNST vs MPOA), * (BNST vs VMHvl) and ‡ MPOA vs VMHvl), unpaired T-tests. **F.** Same as E, but comparing single with double-labelled cell. Clusters are reordered for each so that proportion of each cluster in the single-projecting category is presented as highest to lowest, left to right, to emphasise differences. Differences between single and double-projecting neurons for enriched clusters are indicated in the category that has significantly more enrichment (unpaired T-test). **G.** Summary model of projections from MeA cell types.

**Supplemental Figure 3 – figure supplement 3.**
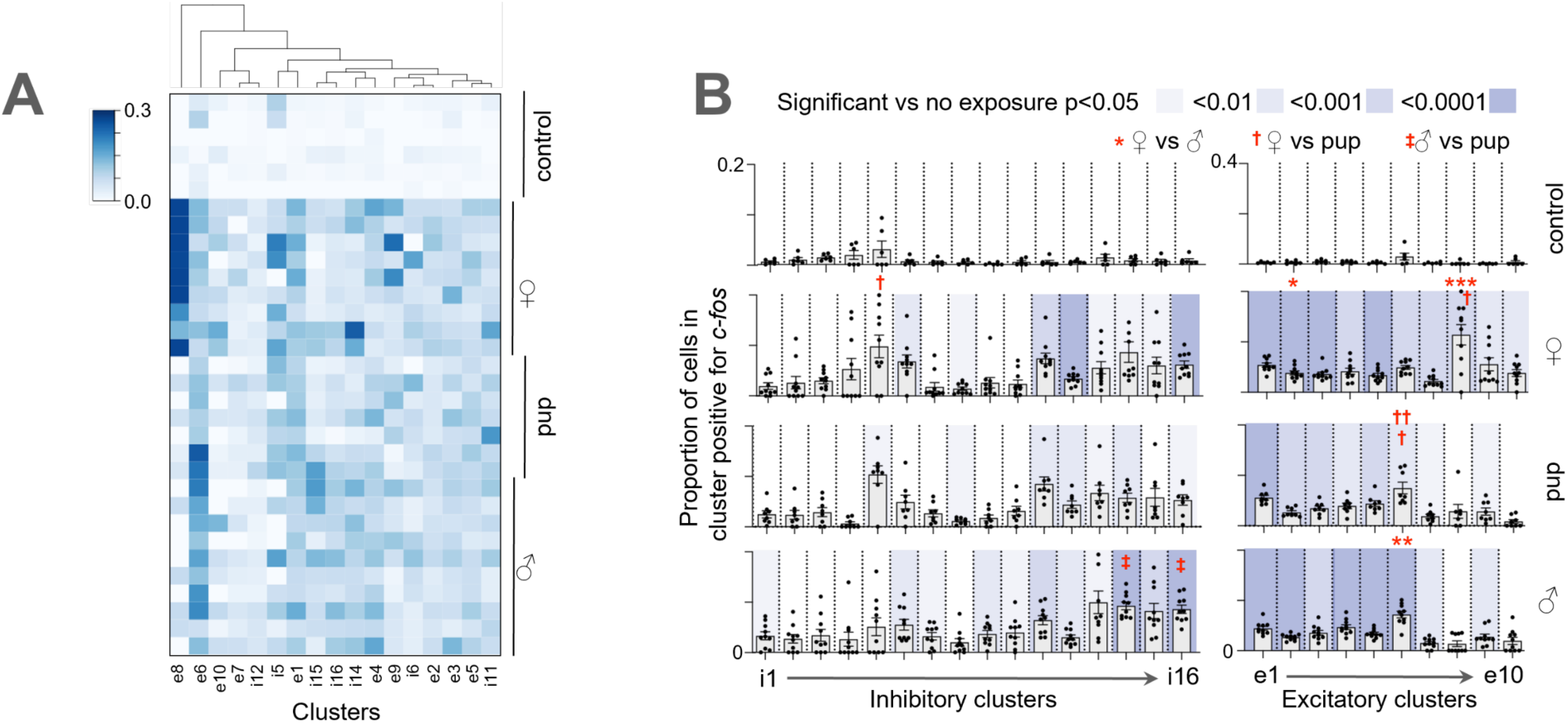
Specific MeA cell types are associated with projection targets and are activated differentially in response to social stimuli. **A.** Heatmap of *c-fos* expression pattern in control, male-exposed, pup-exposed and female-exposed mice. Selected clusters are those with significant responses compared with non-exposed controls. Scale bar represents proportion of cells in cluster positive for *c-fos*. **B.** *c-fos* positive cells per cluster as a proportion of total *c-fos* positive cells (n = 6 (control),10 (female), 8 (pup), 10 (male)). Enrichment indicated by shading (vs non exposure controls), * (female vs male), † (male vs pup), and ‡ (female vs pup), unpaired T-tests.

## References

1 Yao, Z. et al. A high-resolution transcriptomic and spatial atlas of cell types in the whole mouse brain. Nature 624, 317–332 (2023). 10.1038/s41586-023-06812-z

2 Lein, E., Borm, L. E. & Linnarsson, S. The promise of spatial transcriptomics for neuroscience in the era of molecular cell typing. Science 358, 64–69 (2017). 10.1126/science.aan6827

3 Wang, Y. H. et al. EASI-FISH for thick tissue defines lateral hypothalamus spatio-molecular organization. Cell 184, 6361-+ (2021). 10.1016/j.cell.2021.11.024

4 Kishi, J. Y. et al. SABER amplifies FISH: enhanced multiplexed imaging of RNA and DNA in cells and tissues. Nature Methods 16, 533-+ (2019). 10.1038/s41592-019-0404-0

5 Raam, T. & Hong, W. Organization of neural circuits underlying social behavior: A consideration of the medial amygdala. Curr Opin Neurobiol 68, 124–136 (2021). 10.1016/j.conb.2021.02.008

6 Newman, S. W. The medial extended amygdala in male reproductive behavior. A node in the mammalian social behavior network. Ann N Y Acad Sci 877, 242–257 (1999). 10.1111/j.1749-6632.1999.tb09271.x

7 Knoedler, J. R. et al. A functional cellular framework for sex and estrous cycle-dependent gene expression and behavior. Cell 185, 654–671 e622 (2022). 10.1016/j.cell.2021.12.031

8 Chen, P. B. et al. Sexually Dimorphic Control of Parenting Behavior by the Medial Amygdala. Cell 176, 1206–1221 e1218 (2019). 10.1016/j.cell.2019.01.024

9 Player, A. N., Shen, L. P., Kenny, D., Antao, V. P. & Kolberg, J. A. Single-copy gene detection using branched DNA (bDNA) in situ hybridization. J Histochem Cytochem 49, 603–612 (2001). 10.1177/002215540104900507

10 Goh, J. J. L. et al. Highly specific multiplexed RNA imaging in tissues with split-FISH. Nature Methods 17, 689-+ (2020). 10.1038/s41592-020-0858-0

11 Wang, X. et al. Three-dimensional intact-tissue sequencing of single-cell transcriptional states. Science 361 (2018). 10.1126/science.aat5691

12 Hong, W., Kim, D. W. & Anderson, D. J. Antagonistic control of social versus repetitive self-grooming behaviors by separable amygdala neuronal subsets. Cell 158, 1348–1361 (2014). 10.1016/j.cell.2014.07.049

13 Lischinsky, J. E. et al. Transcriptionally defined amygdala subpopulations play distinct roles in innate social behaviors. Nat Neurosci 26, 2131–2146 (2023). 10.1038/s41593-023-01475-5

14 Choi, G. B. et al. Lhx6 delineates a pathway mediating innate reproductive behaviors from the amygdala to the hypothalamus. Neuron 46, 647–660 (2005). 10.1016/j.neuron.2005.04.011

15 Keshavarzi, S., Sullivan, R. K., Ianno, D. J. & Sah, P. Functional properties and projections of neurons in the medial amygdala. J Neurosci 34, 8699–8715 (2014). 10.1523/JNEUROSCI.1176-14.2014

16 Bian, X. Physiological and morphological characterization of GABAergic neurons in the medial amygdala. Brain Res 1509, 8–19 (2013). 10.1016/j.brainres.2013.03.012

17 Abs, E. et al. Learning-Related Plasticity in Dendrite-Targeting Layer 1 Interneurons. Neuron 100, 684–699 e686 (2018). 10.1016/j.neuron.2018.09.001

18 Ibrahim, L. A. et al. Cross-Modality Sharpening of Visual Cortical Processing through Layer-1-Mediated Inhibition and Disinhibition. Neuron 89, 1031–1045 (2016). 10.1016/j.neuron.2016.01.027

19 Schuman, B. et al. Four Unique Interneuron Populations Reside in Neocortical Layer 1. J Neurosci 39, 125–139 (2019). 10.1523/JNEUROSCI.1613-18.2018

20 Canteras, N. S., Simerly, R. B. & Swanson, L. W. Organization of projections from the medial nucleus of the amygdala: a PHAL study in the rat. J Comp Neurol 360, 213–245 (1995). 10.1002/cne.903600203

21 Lin, D. et al. Functional identification of an aggression locus in the mouse hypothalamus. Nature 470, 221–226 (2011). 10.1038/nature09736

22 Chen, K. H., Boettiger, A. N., Moffitt, J. R., Wang, S. & Zhuang, X. RNA imaging. Spatially resolved, highly multiplexed RNA profiling in single cells. Science 348, aaa6090 (2015). 10.1126/science.aaa6090

23 Moffitt, J. R. et al. Molecular, spatial, and functional single-cell profiling of the hypothalamic preoptic region. Science 362 (2018). 10.1126/science.aau5324

24 Zhang, M. et al. Spatially resolved cell atlas of the mouse primary motor cortex by MERFISH. Nature 598, 137-+ (2021). 10.1038/s41586-021-03705-x

25 Shah, S., Lubeck, E., Zhou, W. & Cai, L. In Situ Transcription Profiling of Single Cells Reveals Spatial Organization of Cells in the Mouse Hippocampus. Neuron 92, 342–357 (2016). 10.1016/j.neuron.2016.10.001

26 Shah, S., Lubeck, E., Zhou, W. & Cai, L. seqFISH Accurately Detects Transcripts in Single Cells and Reveals Robust Spatial Organization in the Hippocampus. Neuron 94, 752-+ (2017). 10.1016/j.neuron.2017.05.008

27 Bugeon, S. et al. A transcriptomic axis predicts state modulation of cortical interneurons. Nature 607, 330-+ (2022). 10.1038/s41586-022-04915-7

28 Gyllborg, D. et al. Hybridization-based sequencing (HybISS) for spatially resolved transcriptomics in human and mouse brain tissue. Nucleic Acids Res 48 (2020). ARTN e112 10.1093/nar/gkaa792

29 Sun, Y. C. et al. Integrating barcoded neuroanatomy with spatial transcriptional profiling enables identification of gene correlates of projections. Nat Neurosci 24, 873–885 (2021). 10.1038/s41593-021-00842-4

30 Xu, S. et al. Behavioral state coding by molecularly defined paraventricular hypothalamic cell type ensembles. Science 370 (2020). 10.1126/science.abb2494

31 Choi, H. M. T. et al. Third-generation in situ hybridization chain reaction: multiplexed, quantitative, sensitive, versatile, robust. Development 145 (2018). 10.1242/dev.165753

32 Shah, S. et al. Single-molecule RNA detection at depth by hybridization chain reaction and tissue hydrogel embedding and clearing. Development 143, 2862–2867 (2016). 10.1242/dev.138560

33 Gandin V, K. J., Yang LZ, Lian Y, Kawase T, Hu A, Rokicki K, Fleishman G, Tillberg P, Aguilera Castrejon A, Stringer C, Preibisch S, Zhe. J Liu. Deep-Tissue Spatial Omics: Imaging Whole-Embryo Transcriptomics and Subcellular Structures at High Spatial Resolution. bioRxiv (2024).

34 Kim, D. W. et al. Multimodal Analysis of Cell Types in a Hypothalamic Node Controlling Social Behavior. Cell 179, 713–728 e717 (2019). 10.1016/j.cell.2019.09.020

35 Lischinsky, J. E. et al. Embryonic transcription factor expression in mice predicts medial amygdala neuronal identity and sex-specific responses to innate behavioral cues. Elife 6 (2017). 10.7554/eLife.21012

36 Hirata, T. et al. Identification of distinct telencephalic progenitor pools for neuronal diversity in the amygdala. Nat Neurosci 12, 141–149 (2009). 10.1038/nn.2241

37 Nordman, J. C. et al. Potentiation of Divergent Medial Amygdala Pathways Drives Experience-Dependent Aggression Escalation. J Neurosci 40, 4858–4880 (2020). 10.1523/JNEUROSCI.0370-20.2020

38 Ishii, K. K. et al. A Labeled-Line Neural Circuit for Pheromone-Mediated Sexual Behaviors in Mice. Neuron 95, 123–137 e128 (2017). 10.1016/j.neuron.2017.05.038

39 Ferrero, D. M. et al. A juvenile mouse pheromone inhibits sexual behaviour through the vomeronasal system. Nature 502, 368-+ (2013). 10.1038/nature12579

40 Bergan, J. F., Ben-Shaul, Y. & Dulac, C. Sex-specific processing of social cues in the medial amygdala. Elife 3, e02743 (2014). 10.7554/eLife.02743

41 Li, Y. et al. Neuronal Representation of Social Information in the Medial Amygdala of Awake Behaving Mice. Cell 171, 1176–1190 e1117 (2017). 10.1016/j.cell.2017.10.015

42 Bintu, B. I., Y.; Zhuang, X.; Dulac C. Molecular and Spatial Organization of the Primary Olfactory System and its Responses to Social Odors. bioRxiv 2025.05.02.651832 (2025). 10.1101/2025.05.02.6518 32

43 Xu, Q., Schlabach, M. R., Hannon, G. J. & Elledge, S. J. Design of 240,000 orthogonal 25mer DNA barcode probes. Proc Natl Acad Sci U S A 106, 2289–2294 (2009). 10.1073/pnas.0812506106

44 Yao, R. W., Luan, P. F. & Chen, L. L. An optimized fixation method containing glyoxal and paraformaldehyde for imaging nuclear bodies. RNA 27, 725–733 (2021). 10.1261/rna.078671.120

45 Moffitt, J. R. et al. High-throughput single-cell gene-expression profiling with multiplexed error-robust fluorescence in situ hybridization. Proc Natl Acad Sci U S A 113, 11046–11051 (2016). 10.1073/pnas.1612826113

46 Hao, Y. et al. Integrated analysis of multimodal single-cell data. Cell 184, 3573–3587 e3529 (2021). 10.1016/j.cell.2021.04.048

47 Stringer, C., Wang, T., Michaelos, M. & Pachitariu, M. Cellpose: a generalist algorithm for cellular segmentation. Nat Methods 18, 100–106 (2021). 10.1038/s41592-020-01018-x

48 Babcock, H., Sigal, Y. M. & Zhuang, X. A high-density 3D localization algorithm for stochastic optical reconstruction microscopy. Opt Nanoscopy 1 (2012). 10.1186/2192-2853-1-6

49 Wolf, F. A., Angerer, P. & Theis, F. J. SCANPY: large-scale single-cell gene expression data analysis. Genome Biol 19, 15 (2018). 10.1186/s13059-017-1382-0

50 Mayerich, D., Abbott, L. & McCormick, B. Knife-edge scanning microscopy for imaging and reconstruction of three-dimensional anatomical structures of the mouse brain. J Microsc 231, 134–143 (2008). 10.1111/j.1365-2818.2008.02024.x

51 Ragan, T. et al. Serial two-photon tomography for automated mouse brain imaging. Nature Methods 9, 255–U248 (2012). 10.1038/Nmeth.1854

52 Tyson, A. L. et al. Accurate determination of marker location within whole-brain microscopy images. Sci Rep-Uk 12 (2022). ARTN 867 10.1038/s41598-021-04676-9

53 Niedworok, C. J. et al. aMAP is a validated pipeline for registration and segmentation of high-resolution mouse brain data. Nature Communications 7 (2016). ARTN 11879 10.1038/ncomms11879

54 Tyson, A. L. et al. A deep learning algorithm for 3D cell detection in whole mouse brain image datasets. Plos Comput Biol 17 (2021). ARTN e1009074 10.1371/journal.pcbi.1009074

